# A Pangenomic Method for Establishing a Somatic Variant Detection Resource in HapMap Mixtures

**DOI:** 10.1101/2025.09.29.679336

**Authors:** Nahyun Kong, Zitian Tang, Andrew Ruttenberg, Juan F. Macias-Velasco, Zefan Li, Wenjin Zhang, Benpeng Miao, Zilan Xin, Qichen Fu, Haeorum Park, Xiaoyu Zhuo, Elvisa Mehinovic, Edward Belter, Chad Tomlinson, John E. Garza, Shihua Dong, Emma Casey, Ben Johnson, Mary F Majewski, Theron Palmer, Yuchen Cheng, Tina Lindsay, Tim Schedl, Daofeng Li, Hui Shen, Robert Fulton, SMaHT Network Assembly/Pangenome Working Group, Ting Wang, Sheng Chih Jin

## Abstract

Somatic mosaicism is essential in human biology and disease, yet robust benchmarks are scarce. The SMaHT Consortium mixed six HapMap cell lines to create artificial somatic variants spanning 0.25% to 16.5% variant allele fractions. We developed a technology-agnostic method that builds pangenome graphs from individual assemblies to create unified benchmarking sets: > 6M single-nucleotide variants, 1.8M small insertions/deletions, 49K structural variations, and 10K mobile element insertions across autosomes, X, and mitochondrial chromosomes. We validated the variants using ultra-deep simulated reads and developed a binomial-based model to estimate coverage requirements for variant detection. Evaluating multiple callers showed CHM13 alignment improves structural variant detection and offers advantages in difficult-to-map regions compared to GRCh38. Systematic characterization showed regions with low detection rate are enriched in centromeres, satellite sequences, tandem repeats, and falsely duplicated genes. This accurate, versatile resource enables systematic evaluation of somatic variant detection technologies.

## INTRODUCTION

Somatic variants—genetic alterations that arise in cells after fertilization—are increasingly recognized as key contributors to a wide range of human diseases, including cancer, overgrowth syndromes, neurodevelopmental disorders, and age-related conditions.^1–9^ To advance understanding of the role of somatic variation in health and disease, the Somatic Mosaicism across Human Tissues (SMaHT) Network was launched under the NIH Common Fund (https://smaht.org/).^10^ SMaHT aims to generate a comprehensive reference dataset from 150 donors representing 19 histologically normal tissues and to develop innovative technologies and computational tools that improve somatic variant detection. A key priority of the program is to ensure that the resulting data are readily accessible and interoperable with other large-scale genomic resources to the broader research community.

Accurate identification of somatic variants and reliable assessment of detection methods require well-designed benchmarking resources. Most existing benchmarking sets have been developed in cancer research^11,12^ using matched tumor–normal pairs, while non-cancer studies have primarily relied on parent-offspring trios or large pedigrees.^13,14^ In both contexts, benchmarking set construction typically involves harmonizing and filtering outputs across multiple sequencing technologies and variant callers^11,15–17^. Even when groups employ the same callers, discrepancies often arise due to differences in alignment parameters and criteria for removing low-confidence variants^14^. Furthermore, most published benchmarking sets focus exclusively on single-nucleotide variants (SNVs) and autosomal chromosomes, leaving more complex variants, such as structural variations (SVs; ≥50 bp), mobile element insertions (MEIs), and variants on sex or mitochondrial chromosomes, unevaluated.^14^

To overcome current limitations, we developed a well-characterized sample mixture and an associated benchmarking set that enable systematic evaluation of somatic variant detection (**Figure 1**). As one of the Genomic Characterization Centers (GCCs) within SMaHT, we designed and created the HapMap cell line mixture—a homogeneous mixture of diploid cell lines studied through the HapMap project^18^ and the Human Pangenome Reference Consortium^19^. These cell lines were cultured independently and then pooled at pre-defined ratios to emulate the allelic fractions ranging from 0.25% to 16.5% (**STAR Methods**). The mixture was then profiled with 500× short-read whole-genome sequencing (WGS) and 100× long-read WGS.

**Figure 1.**
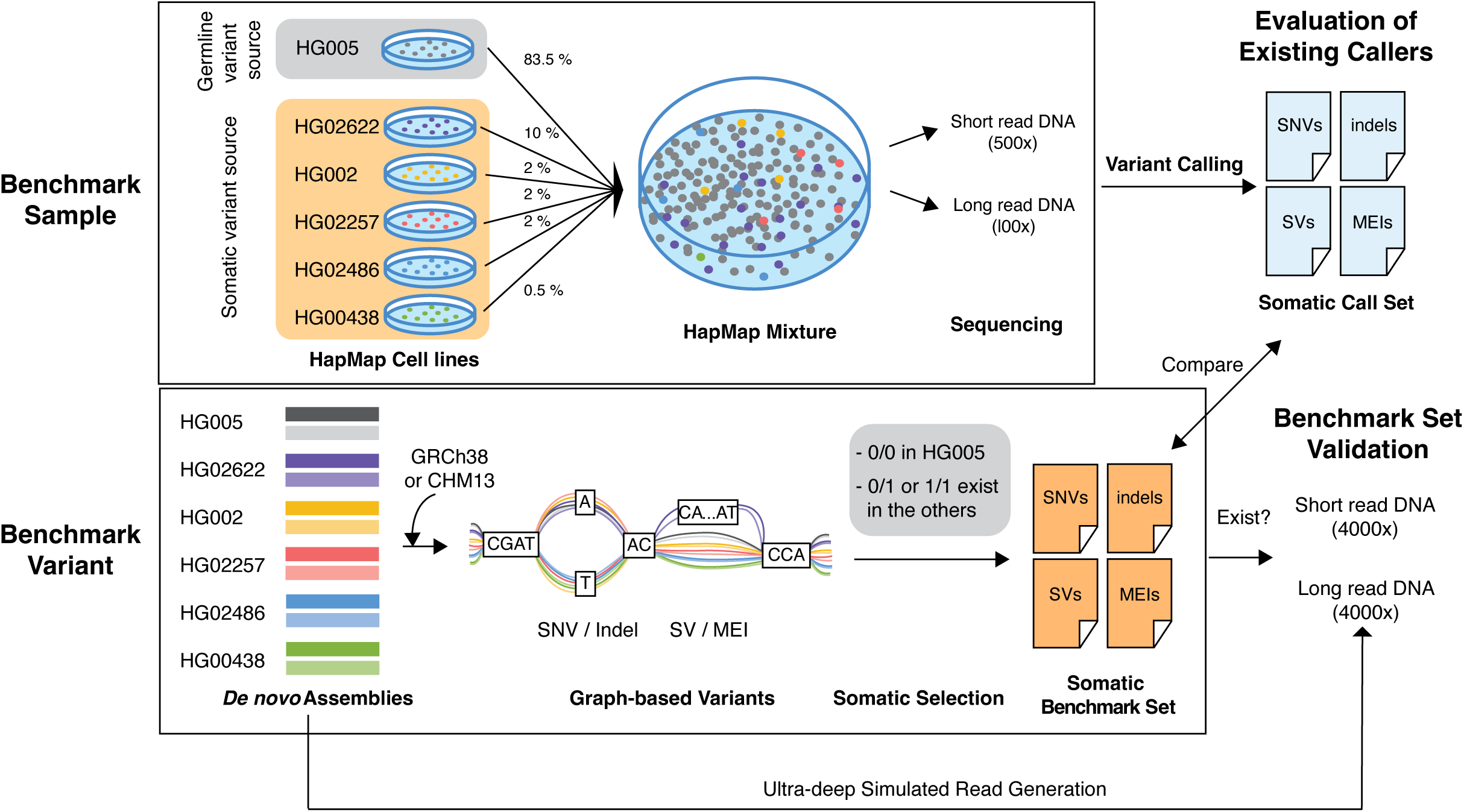
Experimental design for generating and validating the somatic variant benchmarking set. Top: construction of benchmark samples by mixing HapMap cell lines at defined proportions, with HG005 (83.5%) as the artificial germline source, and five other cell lines (10% HG02622, 2% HG002, 2% HG02257, 2% HG02486, and 0.5% HG00438) as artificial somatic variant sources. Bottom: Bioinformatics workflow showing the full process from *de novo* assemblies through MC graph-based variant discovery to final somatic benchmarking set generation. Briefly, maternal and paternal assemblies for all six HapMap samples and a reference (GRCh38 or CHM13) were aligned in parallel, with differences at base-pair resolution in assemblies represented as bubbles. Any variants in the bubbles that were homozygous reference (0/0) in HG005 and homozygous alternative or heterozygous in at least one of the other cell lines were classified as somatic variants for the benchmarking set. Right: This benchmarking set was validated in ultra-deep simulated data and was used to benchmark variants called from HapMap mixture sequencing data. MC, minigraph-cactus; SNVs, single-nucleotide variants; Indels, small insertions and deletions; SVs, structural variants.

To construct the benchmarking variant set, we also developed a novel method using Minigraph Cactus^20^ (MC; **STAR Methods**), integrating both reference and *de novo* assemblies from HapMap samples. This single pangenome graph framework enables the generation of all types of somatic variants—not just in autosomes, but also in the sex and mitochondrial chromosomes—across a wide range of variant allele fractions (VAFs).

Importantly, it is technology-agnostic, remaining robust across different sequencing platforms and adaptable as new technologies and variant-calling methods continue to emerge.

Using this resource, we systematically evaluated the performance of somatic variant callers across mutation types and genomic contexts. We also provided sequencing coverage recommendations for each of 6,637 medically relevant genes, substantiating prior findings that shows that coverage variability is a key determinant of diagnostic yield.^21^ Together, these analyses offer practical guidance to inform clinical sequencing strategies and gene-level sensitivity optimization. More broadly, by providing genome-wide, technology-agnostic benchmarks, our resource overcomes the limitations of relying on rare, benchmarking somatic events in individual samples and provides a powerful framework for detecting, quantifying, and reducing biases in somatic variant analysis across research, commercial, and clinical applications.

## RESULTS

### Design of benchmark sample and data set

To create a resource for comprehensive evaluation of the somatic variants across a broad range of VAFs, six HapMap cell lines were precisely mixed to generate a benchmark sample comprising 83.5% HG005 (Male; Chinese), 10% of HG02622 (Female; Gambian in Western Division, Mandinka), 2% each of HG002 (Male; Ashkenazim Jewish), HG02257 (Female; African Caribbean in Barbados), and HG02486 (Male; African Caribbean in Barbados), and 0.5% of HG00438 (Female; Chinese Han in the South; **Figure 1**), where heterozygous and homozygous variants in HG005 act as germline variant source, and variants in the other five cell lines act as somatic variant sources.^16,17^ HapMap mixture was sequenced to 500× using short-read WGS (Illumina NovaSeq X Plus) and to 100× using long-read WGS (PacBio Revio), yielding high-quality reads for downstream analyses with quality control metrics pre- and post-alignment shown in **Figure S1** and **Table S1-S2**.

### Establishing a benchmark somatic call set and high-confidence genomic regions

To establish a somatic variant benchmarking set, we used *de novo* assemblies from HapMap individuals, alongside GRCh38 or T2T-CHM13 reference assemblies **(Figure 1**). Using GRCh38 and T2T-CHM13 reference as the backbone, respectively, we built an MC graph^20^ to capture the full spectrum of genetic variations—spanning SNVs, short insertions and deletions (indels), and SVs — restricted to benchmark regions, defined as the intersection of reliable regions across all assemblies (**Figure S2**). These regions encompassed 89% of GRCh38 and 90% of CHM13 (**Figure S3**). To define somatic events within the HapMap mixture, we retained only those variants that were homozygous reference in the germline-sourced sample (HG005) and exhibited an alternative genotype in at least one of the somatic-sourced samples (HG02622, HG002, HG02257, HG02486, and HG00438) (see **STAR Methods**). MEI benchmarking sets were generated from the SV benchmarking set variants overlapping with young MEI annotations (**Figure S4f; Figure S5f**).^22^

Using the GRCh38 backbone, the benchmarking set captured over 6 million SNVs, 1.8 million small indels, 51,000 SVs, and 10,000 MEIs (**Table 1**) distributed across all autosomes, chromosome X, and the mitochondrial genome. Both the GRCh38-based and CHM13-based benchmarking sets spanned the expected VAF window (0.25 %–16.5 %) (**Figure S4a-d**) and displayed the expected excess of transitions over transversions (ts/tv = 1.86). SV length distributions exhibited peaks at ∼300 bp and ∼6 kb insertion, consistent with the canonical sizes of Alu and L1 insertions, respectively (**Figure S4e**). Most SNVs, indels, and SVs are unique to a single haplotype (**Figure S4g-j**). We observed similar patterns in the CHM13-based benchmarking set (**Figure S5**). Moreover, we found a high proportion of nested variants—defined as variants that overlap with other variants—for SVs (80% of all SVs in our benchmark set), with 3% of SNVs and 58% of indels classified as nested.

**Table 1.** Summary of benchmarking set variant counts.

| Reference version | Covered Region |  | Variant Count |  |  |  |
| --- | --- | --- | --- | --- | --- | --- |
|  | Base (bp) | % genome | SNV | Indel | SV | Total |
| GRCh38 | 2,751,815,713 | 89.1% | 6,037,703 | 1,832,989 | 51,006 (including 10,607 MEIs) | 7,921,698 |
| CHM13 | 2,788,359,512 | 89.4% | 6,443,434 | 2,025,509 | 49,109 (including 9,417 MEIs) | 8,518,052 |

Genomic distribution analysis revealed that these nested SVs cluster within known SV hotspots, previously identified by the Human Genome Structural Variation Consortium (**Figure S6**).^23^

### Validation demonstrates the reliability of the benchmarking set

To assess the accuracy of the benchmarking set, we performed validation using error-free, ultra-deep simulated sequencing data generated from each HapMap assembly, preserving the exact mixture proportions used in the actual HapMap mixture (**Table S1)**. Our simulated short-read and long-read datasets closely resembled the original sequencing data in GC content and other quality metrics **(Table S1 and Figure S1)**.

To determine the optimal coverage for validation, we simulated short reads from 500× to 4500× in 500× increments. Validation rates increased with coverage and plateaued at 4000×, changing by less than 0.1% thereafter (**Figure S7a**). At 4000×, recall values were consistent across different VAFs, yielding an overall SNV validation rate of 0.956 – 0.991 (**Figure S7b**). We thus selected 4000× for subsequent analyses.

At simulated 4000×, we validated SNVs and indels using short reads and SVs using long reads. A variant is considered validated if supported by at least one read carrying the alternative allele. SNVs and indels required an exact coordinate-and-allele match, and SVs were assessed with Truvari (default 70% threshold for both sequence and size similarity).^24^ Using alignment-based approaches, validation rates for SNVs, indels, and SVs were 0.933, 0.867, and 0.937, respectively (**Figure 2a**). Close examination of unvalidated variants revealed that most were present in the simulated reads but missed either due to misalignment (**Figure S8a**) or differences in representation between graph and linear genome alignment methods (**Figure S8b**). By incorporating assembly-based validation approaches (see **STAR Methods**), where we validated the sequence context present in the source *de novo* assembly, we boosted the validation rates to 94.0% for SNVs, 96.8% for indels, and 96.0% for SVs (**Figure 2a**).

**Figure 2.**
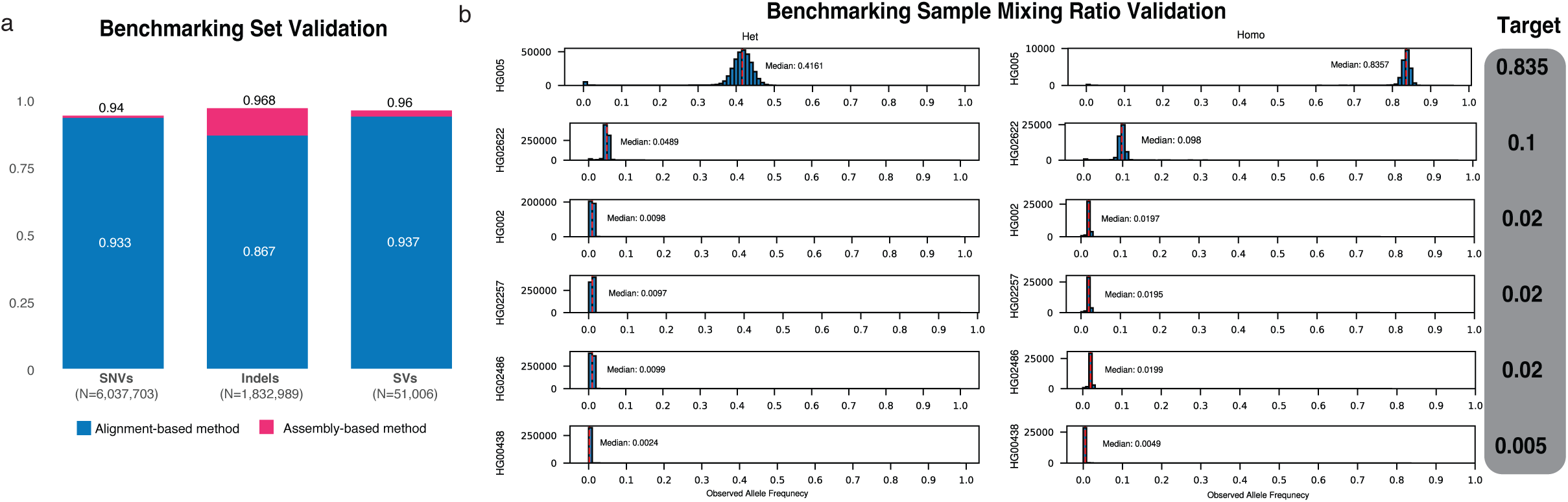
Benchmarking set validation rate and HapMap mixing ratio check using benchmarking set. (a) Variant validation rates through alignment-based (blue) and assembly-based (pink) methods for different variant types: SNVs, indels, and SVs that include MEIs. (b) VAF distribution of heterozygous or homozygous variants exclusively in one of the six HapMap samples, as measured in real 500× data. The median VAF of each plot (labeled by red dotted line) estimates the mixing ratio of that cell line (if homozygous) or half the mixing ratio (if heterozygous). VAF, variant allele frequency.

Most unvalidated variants were in inherently difficult regions. For instance, of all SNVs missed in alignment-based validation on chromosome 1, 73.7% were in genomic distortion regions. These regions are defined as such that reads can easily fail to map back to where they are derived from, thereby affecting alignment (see **STAR Methods**). On the other hand, 74% of SNVs missed by assembly-based validation were in Dipcall non-confident regions.^25^ Overall, of variants failing both alignment- and assembly-based validation, 95.2% of SNVs, 70.5% of indels, and 62.4% of SVs fell within either distortion or Dipcall non-confident regions (**Table S3**), underscoring the challenge of these loci and supporting the reliability of the benchmark elsewhere.

### Validation of HapMap sample mixture ratio

Having established the reliability of our benchmark, we next validated whether the HapMap cell lines are mixed in the intended ratio. Using the WashU 500× short-read sequencing data, we compared benchmark SNVs with variants observed in the sequencing data. For each sample’s unique heterozygous or homozygous SNVs in the benchmark, we computed the observed VAFs, visualized their distribution with frequency histograms, and recorded the median of these VAF values as our primary metric for assessing the mixing ratio. Overall, the mixing ratio was consistent with the expected ratios (**Figure 2b**).

We next evaluated how accurately aligned data reflects VAFs for the SNVs spanning a broader VAF range. Using the 4000× simulated short-read data, we compared each variant’s observed VAF with its expected value from the mixture design. Most SNVs exhibited minimal deviation (**Figure S9a**). We further quantified a variant allele count range based on a binomial distribution using its expected VAF and the total read depth at that locus, and considered the observed allele count to be justified if it fell within the 95% confidence interval (CI). Using this approach, 96.8% of our benchmark SNVs were within the 95% CI.

We then investigated the remaining 3.2% of SNVs whose observed VAFs deviated from the expected value beyond what the random sampling can account for. Using the previously described binomial distribution, we defined a null hypothesis as the expected VAF being equal to the observed VAF, and the alternate hypothesis as a significant difference between the expected and observed VAFs (see **STAR Methods**). Interestingly, there was a marked enrichment of somatic events in difficult-to-map regions and regions with high distortion metrics (Pearson’s χ² = 128,283 and 153,282, respectively; p < 2.2 × 10LJ¹LJ for both tests; **Figure S9b**; see **STAR Methods**). These distortions may occur when the reference is insufficiently representative or when the complex sequence structure confounds the aligner. Taken together—and considering that VAF fluctuations are likely to be further amplified at typical sequencing depth (100∼500×)— our results indicate that, while most somatic SNVs can be captured at expected VAF, variants arising from repetitive or structurally complex regions are susceptible to VAF distortion.

### Modeling a theoretical framework for variant detection rates across sequencing depths

Determining the optimal sequencing depth for reliably detecting somatic variants has long been a challenging question in genomics. To estimate the sequencing depth required to detect somatic variants, we measured the percentage of benchmarking set variants supported by ≥1 alternate read using short-read sequencing data generated by four different SMaHT GCCs, with their sequencing depths ranging from 171× to 491× (**Figure 3a**). As anticipated, detection rates increased with both deeper coverage and higher VAF. To generalize this observation, we modeled the detection rate with simple binomial sampling, treating every read as an independent draw with success probability equal to the variant’s VAF. This baseline captured the overall trend but systematically overLJpredicted detection above ∼2% VAF and underLJpredicted detection below ∼0.5% VAF. The highLJVAF deficit can be explained by sequencing or aligner errors that remove true positive alternate reads, whereas the surplus at very low VAF may stem from sequencing noise that converts reference reads into apparent alternates (**Figure S10a-b**).

**Figure 3.**
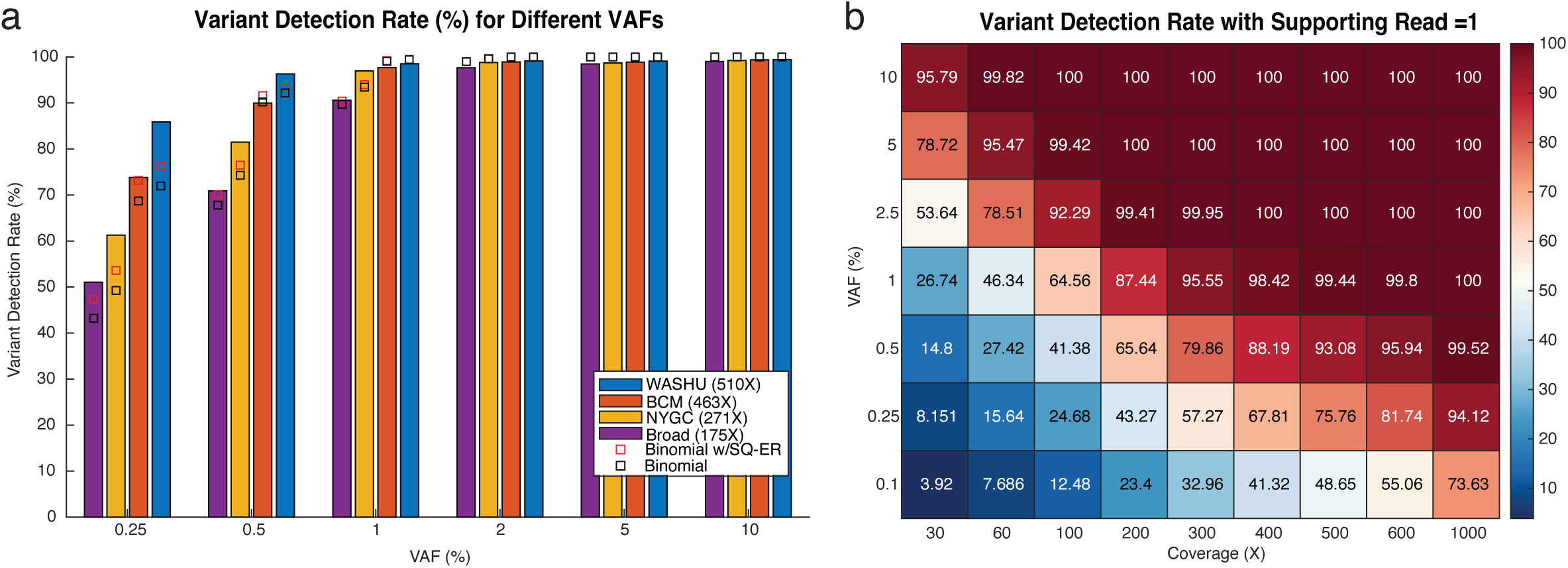
Optimal sequencing depth for variant detection at different VAFs. (a) SNVs detection rates measured at varying VAFs based on sequencing data generated at four different sequencing centers (WashU: Washington University in St. Louis, NYGC: New York Genome Center, BCM: Baylor College of Medicine, and Broad: Broad Institute), with binomial model predictions with (red) and without (black) sequencing error shown in squares. Here, the detection rate was defined as observing at least one supporting read for the variant in the raw BAM file. (b) Heatmap showing theoretical detection rate of variants at different sequencing depths and VAFs. Variants are considered “detected” if they appear in at least one read (k0=1). SNVs, single-nucleotide variants; VAF, variant allele frequency.

To improve the fit, we developed two additional statistical models that could incorporate sequencing error (see **STAR Methods**). The best fit was obtained with a binomial sampling model augmented by a binomial sequencing-error component, which achieved the lowest root-mean-square deviation and highest correlation with the empirical detection rates (**Figure 3a**; **Figure S10**).^26^ This prediction model provides a practical guide for determining the coverage requirements needed to reach a maximum sensitivity for variants at specific VAF thresholds: for instance, ∼300× coverage is required to capture >95% of SNVs present at 1% VAF (**Figure 3b**). We also extended the model to incorporate varying threshold for the number of supporting reads and provided the corresponding metrics (**Figure S11**). As expected, higher coverage is required when more supporting reads are needed to detect a variant call.

Together, these analyses provide a theoretical framework for planning sequencing depth to achieve desired sensitivity across VAF ranges. While each somatic variant caller imposes its own probabilistic models and coverage-dependent filters^25,27,28^, our framework sets an upper bound on achievable sensitivity, offering a reference point for developers and users when benchmarking caller performance and guiding sequencing study design.

### Somatic variant calling performance comparison

We next evaluated variant caller performance across VAF strata using our benchmark (**Figure 4a**). For 500× short-read WGS, Strelka2^29^ achieved the highest precision (98% for GRCh38 SNVs; 92% for indels) but with moderate sensitivity. VarScan2^30^ delivered the best sensitivity (84% for SNVs; 39% for indels) at the expense of precision. Mutect2^27^ struck the best balance, with 89% precision and 68% sensitivity for SNVs, and 77% precision with 29% sensitivity for indels.

**Figure 4.**
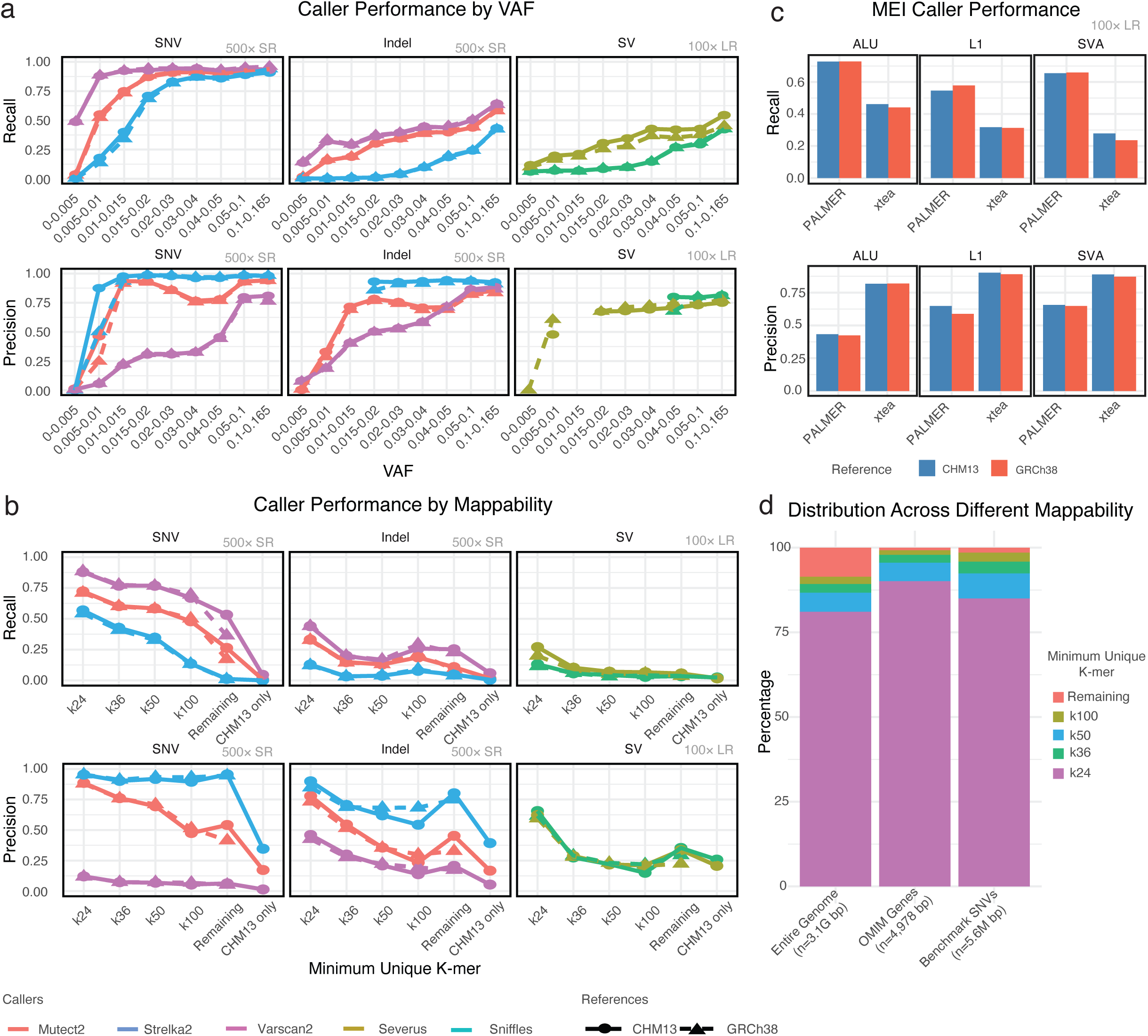
Somatic variant caller evaluations. (a) Recall and precision plots for six different somatic variant callers (Mutect2, Strelka2, Varscan2, Severus, Sniffles, xTEA, and PALMER) measured with variants across various VAF bins at short read (SR) and long read (LR) real data. Both recall and precision drop as VAF decreases. (b) Recall and precision plots for six different somatic variant callers (Mutect2, Strelka2, Varscan2, Severus, Sniffles, xTEA, and PALMER) in regions of different mappability stratified by UMAP k-mers. (c) Recall and precision plots for two MEI callers (xTea and PALMER) with different MEI types (ALU, L1, and SVA). (d) Percentage base-pairs distribution across various mappability regions for the entire human genome, OMIM genes, and the benchmarking set SNVs. VAF, variant allele frequency; indels, small insertions and deletions; MEI, mobile element insertion; SNV, single-nucleotide variant.

For 100× long-read WGS, Severus^31^ achieved higher recall than Sniffles2^28^; however, because Sniffles2 did not detect SVs at reported VAFs below 5%, relative precision could not be assessed in this lowLJVAF range. This finding underscores the need for deeper coverage to reliably capture low-VAF SVs. Furthermore, unlike other variant types, SV caller performance varied depending on the reference genome. On average, precision was higher on GRCh38 (75.5% vs. 74.4%), whereas CHM13 yielded greater sensitivity (21.2% vs. 18.5%) and a modestly improved F1 score (0.323 vs. 0.293) (**Figure S12**). For MEIs from 100× longLJread data, PALMER^32^ outperformed xTea^33^ in longLJread mode (**Figure 4c**).

PALMER’s repeatLJaware preLJmasking and kLJmer scanning strategy retrieves MEI fragments hidden within other repeats, whereas xTea—originally optimized for short reads— depends chiefly on split/full alignments and therefore misses such insertions.

To evaluate the influence of sequence context, we stratified results by UMAP kLJmer mappability^34^, a measure of how uniquely sequencing reads of a given length (k) align to the reference genome (**Figure 4b**). Sensitivity of all callers drops as mappability decreased, with the highest performance in easy-to-map regions (k24) and lower performance in difficult-to-map regions (k36–k100 and the remaining regions), while precision changed little. In k24 easy-to-map regions, alignment to CHM13 yielded slightly higher recall and precision than GRCh38 across all variant classes (average F1: 0.573 vs. 0.562 for SNVs, 0.379 vs. 0.367 for indels, and 0.298 vs. 0.247 for SVs; **Figure 4b; Figure S12**). In the most difficult-to-map regions, CHM13 alignments again demonstrated superior performance, where even 100-mers cannot be uniquely placed (remaining regions), particularly for SNVs and indels (**Figure S12**; average F1: 0.164 vs. 0.117 for SNVs and 0.164 vs. 0.137 for indels, 0.080 vs. 0.063 for SVs compared to hg38; **Figure 4b; Figure S12**). In contrast, regions unique to CHM13 yielded the lowest performance, reflecting their sequence complexity (average F1: 0.013 for SNVs, 0.032 for indels, 0.038 for SVs).

Although ∼81% of the genome lies in k24 easyLJtoLJmap sequence, ∼9% of regions of disease-causing genes reside in k36-k100, or more challenging intervals (**Figure 4d**), making them clinically crucial yet analytically demanding. Notably, 16% of variants in our benchmark also fall in these regions, making the dataset as an invaluable resource for improving somatic variant detection in low-mappability regions.

### Characterization of genomic regions with failed somatic variant detection

To understand regional biases in somatic variant calling, we investigated genomic loci where callers failed to detect variants. We focused on SVs given that their detection is complicated by structural heterogeneity and breakpoint ambiguity. For this analysis, the genome was partitioned into 1 kb windows and classified as either SV-harboring regions (containing benchmark SVs) or regions with no true positive calls (noTP regions) (**Figure 5a**; see **STAR Methods**).

**Figure 5.**
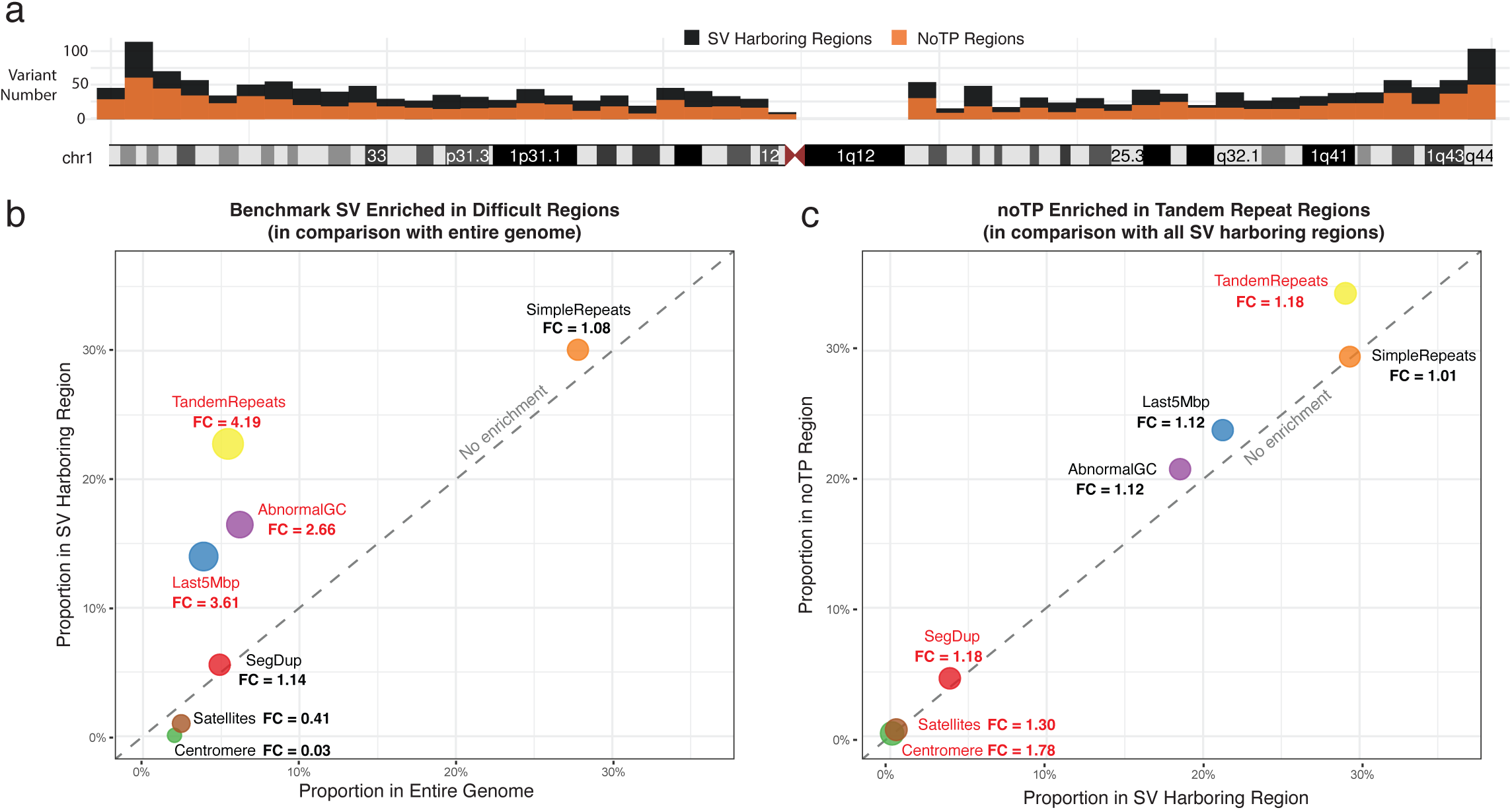
Genomic distribution of structural variant detection failures across challenging sequence contexts. (a) Schematic representation of SV harboring regions (black bars) and regions with no true positive calls (noTP regions, orange bars) across chromosome 1. SV harboring regions contain benchmark structural variants, while noTP regions represent genomic windows where variant callers failed to detect any true positive variants. (b) Enrichment analysis of benchmark structural variants in difficult genomic regions compared to genome-wide distributions. Each point represents a different challenging sequence feature, with fold change (FC) values indicating enrichment relative to the entire genome. Points above the diagonal dashed line (no enrichment) indicate overrepresentation in SV harboring regions. Tandem repeats show the highest enrichment (FC = 4.19), followed by subtelomeric regions within the last 5 Mbp of chromosomes (FC = 3.61) and regions with abnormal GC content (FC = 2.66). (c) Enrichment analysis of detection failures (noTP regions) within SV harboring regions after restricting analysis to variants with ≥10% VAF. Comparison shows modest enrichment of failed calls in centromeric regions (FC = 1.78), satellite sequences (FC = 1.30), tandem repeats (FC = 1.18), and segmental duplications (FC = 1.18). Point size reflects the relative fold enrichment of each genomic feature.

We first quantified the proportion of challenging genomic features^35^ across the entire genome, SV-harboring regions, and noTP regions. Benchmark SVs showed significant enrichment in tandem repeats (fold change [FC] = 4.19), subtelomeric regions within the last 5 Mbp of each chromosome (FC = 3.61), and regions with abnormal GC content (FC = 2.66), compared to genome-wide random expectations (**Figure 5b**). However, comparing noTP regions against SV harboring regions revealed no substantial enrichment in any specific difficult genomic feature (**Figure S13**), indicating that caller failures occur broadly across various challenging regions rather than concentrating in a single genomic context.

Since variant calling was performed on 100× long-read data, we next asked whether many noTP regions resulted from insufficient local coverage rather than inherent sequence difficulty. To mitigate the effect of coverage, we restricted our analysis to SV harboring regions containing benchmark variants with ≥ 10% VAF and redefined the subset lacking TP calls as the refined noTP set. Under this more stringent definition, modest enrichment emerged in several difficult contexts, including centromeric regions (FC = 1.78), satellite sequences (FC = 1.30), tandem repeats (FC = 1.18), and segmental duplications (FC = 1.18) (**Figure 5c**). These findings indicate that both sequence complexity and sequencing coverage limit somatic SV calling performance, with low-frequency variants being particularly susceptible.

### Coverage recommendations for detecting somatic variants in clinically relevant genes

Somatic variants are increasingly recognized as important contributors to cancer and other diseases, and their detection has become an essential component of clinical genetic testing.^3,7,36,37^ As a result, clinical laboratories are under growing pressure to reliably detect low-VAF somatic variants using high-depth sequencing technologies. A persistent additional challenge, however, lies in uneven genome coverage. Differences in GC content, repetitive elements, and local mappability lead to systematic underrepresentation of certain regions, including those of high clinical relevance.^21^

To address this gap, we compiled an updated list of 6,633 clinically relevant genes located on the autosomes and chromosome X (**Table S5**). This set expands on a previously curated group of 4,697 genes^38^ by integrating multiple major sources. Specifically, we included 4,970 genes from the Online Mendelian Inheritance in Man^39^ (accessed December 29, 2024), 84 actionable genes from the American College of Medical Genetics and Genomics SF v3.3 list^40^, 736 cancer-related genes from the COSMIC Cancer Gene Census^41^ (accessed March 2023), and 83 genes implicated in clonal hematopoiesis from recent reports.^42–44^

We next modeled the relationship between sequencing coverage and somatic variant detection performance using benchmark SNVs at 1% VAF (N = 414,309) located within the 6,633 clinically relevant genes (see **STAR Methods**). For this analysis, we used Mutect2— the SNV caller that achieved the highest F1 score—to measure per-gene variant detection rate across short-read datasets sequenced from Washington University in St. Louis (WashU; 508×), New York Genome Center (NYGC; 271×), Baylor College of Medicine (BCM; 463×), and Broad Institute (Broad; 175×). Using a multivariate exponential model, we estimated the minimum sequencing depth required to capture 75%, 80%, 90% and 95% of variants at 1% VAF (**Table S6; STAR Methods**). Among the 4,612 genes with ≥10 benchmark SNVs, the mean predicted Mutect2 detection rate at 30× was 6.04%, highlighting potential limitations in low-frequency variant detection at conventional 30× sequencing depths. It was also substantially lower than the theoretical 26.74% capture rate predicted by our binomial model for 1% VAF variants with a single supporting read (**Figure 3b** and **6b**), underscoring the difficulties for variant callers to confidently capture all existing variants in the real world. Of these 4,612 genes, 4,564 genes (98.9%) required > 500× coverage to capture 80% of these variants (**Figure 6b**), with *BCL2L1-AS1* requiring the lowest coverage (371×) and *UNC93A* requiring the highest (51,780×) (**Figure 6c**). While such high coverages present practical challenges for whole-genome clinical sequencing, they are potentially achievable through targeted or panel sequencing.

**Figure 6.**
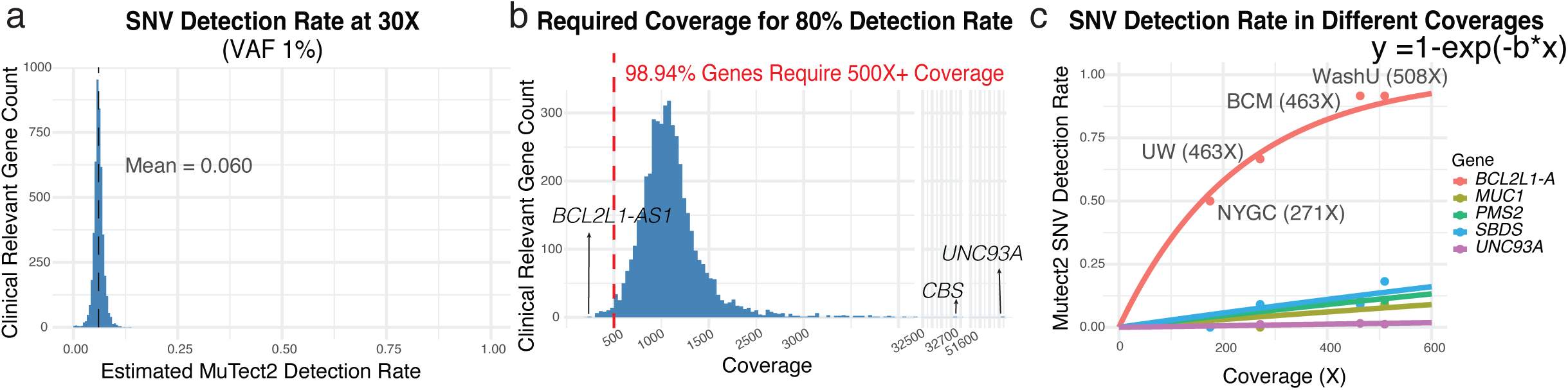
Gene-specific coverage requirements for somatic variant detection at 1% VAF. (a) Distribution of estimated variant detection rate at 30× cross 4,473 clinically relevant genes with ≥10 benchmark SNVs at 1% VAF. Coverage requirements were estimated using a multivariate exponential model applied to Mutect2 performance measured at four different sequencing coverages. (b) Distribution of the minimum sequencing coverage required to capture 80% of somatic SNVs with 1% VAF. *BCL2L1-AS1* required the lowest coverage, while *UNC94A* required the highest. Most genes (98.94%) required minimum of 500× coverage to achieve the 80% sensitivity threshold. (c) Exponential curves showing variant detection rates across coverage depths for representative clinical genes. Observed detection rates at four sequencing depths (510×, 463×, 271×, 175×) were fitted to the exponential model LJ = . Model parameter, *b*, varied substantially between genes, reflecting differences in sequence complexity, mappability, and broader genomic context. *BCL2L1-AS1* showed rapid saturation at relatively low coverage (*b* = 4.34×10^−3^), whereas genes such as *MUC1, PMS2, SBDS,* and *UNC93A* exhibited poor detectability even at high coverage depths (*b* = 1.57 ×10^−4^, 2.37×10^−4^, 2.92×10^−4^, and 3.11×10^−5^, respectively) . SNV, single-nucleotide variant; VAF, variant allele frequency.

Crucially, a small subset of genes (N = 28)—including *SBDS, MUC1,* and *PMS2* —required extremely high coverage (≥2,000×) to achieve an 80% detection rate in this model (**Figure 6c**). Notably, ten of these genes—*CBS, SIGLEC16, MUC1, CRYAA, IFITM3, GTF2IRD2, HBG1, KCNE1*, and *NPPA*— fall within the previously curated set of 273 challenging and medically relevant genes^38^ . This included false-duplication genes in GRCh38 (e.g., *CBS, CRYAA, KCNE1*), where spurious duplicated sequence causes Illumina reads from one haplotype to mis-map to the other copy^38^. Likewise, the reduced detectability in these genes may be largely attributable to sequence complexity, such as low mappability, high GC content, or alignment difficulties within segmental duplications.

Rather than recommending a uniform sequencing depth across all genes, our analysis provides per-gene coverage benchmarks tailored to a specific VAF threshold. These recommendations, provided in **Supplementary Table 6**, enable clinicians and researchers to optimize sequencing strategies for their diagnostic panels or research questions. By linking gene-level coverage requirements with expected detection sensitivity, this resource offers practical guidance for more accurate, efficient, and cost-effective somatic variant detection in both clinical and translational settings.

## DISCUSSIONS

We present a graph-based framework to create a somatic variant benchmarking set. Unlike traditional approaches that rely on aggregating caller outputs, the MC graph we built explicitly captures every possible variant from individual assemblies within the benchmarking region and subjects all variant classes (SNVs, indels, SVs, MEIs) to a single, consistent pipeline. This strategy eliminates sequencing- and caller-specific biases and obviates the need for an external “true negative” set when evaluating precision. Moreover, this approach is coordinate-flexible: we can readily build benchmark sets in any coordinate system by substituting the backbone reference assembly used in the graph pipeline^20^. Here, we generated two benchmark sets on both T2T-CHM13 and GRCh38 and used them to compare somatic-caller performances across references. Looking ahead, using donor-specific assemblies as reference coordinates to construct benchmarks could further reduce germline confounding—effectively removing germline variants from consideration—and improve the precision and recall of somatic variant detection.

Using this framework, we generated the first graph-based somatic variant benchmark from a HapMap cell-line mixture engineered to span pre-defined VAF ranges. This design addresses key limitations of previous benchmarks, which often relied on consolidating unknown or caller-reported VAFs from experimental sequencing data that may deviate significantly from the actual variant frequency.^45^ For example, at 500× coverage, variants with a true 2% VAF can exhibit observed frequencies ranging from 0.96% to 3.6% under a binomial model at 95% CI, due to inherent random sampling in genome sequencing. Our mixture design allowed us to precisely control VAF distributions, ensuring that observed frequencies aligned with expectations and thereby providing a robust framework for benchmarking low-frequency variants.

To our knowledge, this is the first somatic benchmark that spans not only all autosomes but also the mitochondrial genome and X chromosome. Inclusion of mitochondrial DNA is particularly valuable because of its unique biology: high copy number, rapid mutation rate, maternal inheritance, and tissue-specific heteroplasmy.^46–48^ mtDNA variants may be present in all molecules (homoplasmy) or in a fraction (heteroplasmy), and heteroplasmy levels can drift with age and across tissues, contributing to diverse human diseases. Short-read analyses are further complicated by nuclear mitochondrial DNA segments (NUMTs), which can mislead alignment and inflate apparent heteroplasmy. Recent mtDNA callers such as Himito (manuscript in preparation) have already leveraged our HapMap mixture to benchmark mitochondrial variant detection. Similarly, benchmarking on the X chromosome addresses sex-specific challenges, including hemizygosity in males, which alters expected VAFs and demands sex-aware evaluation of callers.^49^

We also developed two statistical models for precisely predicting the required sequencing depth for detecting various somatic variants across different VAF thresholds. The first model estimates the maximum detection rate across the genome by measuring the probability of observing *k* supporting reads for a given variant. This model incorporates sequencing error rates and can be adapted to different sequencing platforms, making it useful for developers and tool designers exploring a wide range of expected VAFs. The second model, a direct extension of the first, captures the actual detection rates achieved for each gene using the Mutect2 SNV caller. These approaches enable more efficient allocation of sequencing resources to address critical clinical and biological questions. In cancer genomics research, for instance, advanced tumors often harbor higher-frequency variants (>5∼10%), while localized tumors may involve ultra-low-frequency variants (<1%)^50^. This underscores the need to customize sequencing strategies depending on experimental settings.

Our evaluation of somatic variant calling performance based on the benchmark set reveals substantial differences in detection capabilities across variant types, sequencing platforms, and genomic contexts. Performance degrades predictably in repetitive regions, with callers maintaining precision while missing true variants in challenging genomic contexts. The regional analysis of variant calling failures provides crucial insights into the genomic determinants of detection performance. Benchmark SVs show pronounced enrichment in challenging sequence contexts—particularly tandem repeats (FC = 4.19) and subtelomeric regions (FC = 3.61). This enrichment suggests that sequence complexity contributes to caller limitations in difficult genomic regions. Furthermore, the modest enrichment observed in regions lacking true positive calls after VAF filtering indicates that coverage limitations also impact somatic variant detection performance. These findings underscore the need for increased sequencing depth and algorithmic improvements to reliably detect low-frequency variants across diverse genomic contexts, particularly in the ∼16% of variants falling within challenging mappability regions that remain analytically demanding yet clinically crucial.

We anticipate that the SMaHT Network and the broader research community will leverage these advances to develop better tools and to generate more accurate somatic—and germline—variant calls, opening the door to answering key questions: (i) Do somatic variants in healthy tissues arise through mutational processes distinct from those that shape germline variants? (ii) To what extent do different healthy tissues share recurrent variants, and what mechanisms—such as mutational hotspots, technical artifacts, or convergent evolutionary pressures—drive their emergence? Addressing these questions will deepen our understanding of genome stability across human tissues and guide future precision-medicine strategies.

## Limitations of the study

Despite these advances, our approach has limitations. First, somatic variants in our benchmarking set were artificially created by mixing the germline variants from HapMap samples. Because germline and bona fide somatic mutations have different mutational signatures and context dependencies, the signals we observe may not fully capture the biology of true somatic events.^51^

Second, our benchmarking set was derived from *de novo* assemblies that represent individual germline variants. As a result, any true somatic mutations originating from the germline variant source sample (HG005) are absent from our benchmarking set and would be labeled false positives.

Third, we observed differences in how nested variants are represented in the graph-based benchmarking set versus the traditional linear reference genome (**Figure S13**). This discrepancy likely stems from graphs constructing SVs prior to any embedded indels or SNVs, whereas linear genome alignments capture each variant independently. Through extensive effort, we reconstructed the nearby sequences and validated these variants, but the discrepancy highlights the limitations of evaluating linear-genome-based calls against a graph-constructed benchmark. Future work should prioritize graph-based callers or robust projection methods to ensure consistency with the graph-based benchmarking set.

Fourth, although our benchmarking set spans all autosomes, chromosome X, and mitochondrial genome, we did not generate a chromosome-Y benchmark because available Y assemblies are haploid and only present in HG002, HG005, and HG02486.

Finally, our MEI benchmark was generated by selecting insertion events that overlap known MEI sequences within the SV benchmark; consequently, this approach may inadvertently miss uncharacterized MEIs absent in current MEI libraries or could inadvertently include inactive MEIs. Addressing these issues will further align the benchmark with biologically relevant somatic variation and improve assessments of low-frequency variant detection.

## RESOURCE AVAILABILITY

### Lead contact

Further requests for information and resources should be directed to and will be fulfilled by the lead contact, Sheng Chih Jin.

### Materials availability

This study did not generate new reagents.

### Data and code availability

The HapMap mixture datasets generated through the SMaHT Project are publicly available at https://data.smaht.org. The data for the patient samples cannot be shared under the restrictions placed by the institutional review board. The benchmarking somatic variant datasets and associated analysis code are available at https://github.com/jinlab-washu.

The final benchmarking set VCF files for SNVs, indels, and SVs is available as separate resources https://wangcluster.wustl.edu/~juanfmacias/Graph_based_HapMap_Truth_Set/.

The graph-based benchmarking set generation pipeline is provided at https://github.com/ztang99/Graph-Based-Benchmark. The scripts for running all analyses and validations, as well as generating all figures included in this manuscript are provided at https://github.com/jinlab-washu/HapMap-BenchmarkSet-Manuscript. The optimal coverage calculator is provided at https://github.com/NahyunKong/Sequencing-coverage-calculator.

## ACKNOWLEDGEMENTS

We thank Genome Technology Access Center at McDonnell Genome Institute at Washington University School of Medicine for sequencing data generation and management. We also thank all members in the Somatic Mosaicism across Human Tissues Network. We thank Eric Roberts in the Hoffman lab at the University Health Network for the Princess Margaret Cancer Centre for providing Uniform Manifold Approximation and Projection K-mer region files in CHM13 coordinates. These studies were supported through NIH grants UM1DA058219, 3UM1DA058219-01S1, U24NS132103, U41HG010972, and R35GM152192.

## AUTHOR CONTRIBUTIONS

N.K., Z.T., R.F., T.W., and S.C.J. contributed to study design and conceptualization. R.F., B.J., M.F.M, and T.P. performed cell culture and sequencing data production. T.L. managed sequencing data producing pipeline. C.T. and D.L. managed sequencing data transfer. N.K., Z.T., A.R., J.F.M., B.M., E.B., C.T., J.E.G., W.Z., Z.X., Q.F., E.M., H.P., X.Z., Z.L., S.D., E.C., Y.C. performed bioinformatics analyses. N.K., Z.T., and A.R. performed statistical analyses. N.K., Z.T., A.R., J.F.M., Z.L, H.P., and W.Z. generated figures. N.K., Z.T., A.R., and S.C.J. wrote the initial manuscript. T.W., R.F., H.S. J.F.M., T.S., and S.C.J. reviewed and edited the manuscript. S.C.J., T.W., R.F., and H.S. administered the project. S.C.J., T.W., B.F., and H.S. provided funding and resources.

## STAR*METHODS

Detailed methods are provided in the online version of this paper and include the following:

### EXPERIMENTAL MODEL AND STUDY PARTICIPANT DETIALS

#### Design and generation of the HapMap mixture

Due to the variable cell quantity requirements across different cell lines for the HapMap design protocol, recovery was staggered to ensure simultaneous harvest time. Each cell line was recovered and expanded individually before being pooled together at designated concentrations immediately prior to cryopreservationon. A desired mix ratio of 83.5% HG005, 10% HG02622, 2% of each of HG002, HG02257, and HG02486, and 0.5% HG00438 was the target for the pool.

Each cell line was thawed and placed in T25 flasks containing 10 ml culture medium (84% RPMI-1640, 15% FBS, and 1% Glutamax). A cell count was removed from the suspension, and the culture volume was adjusted as necessary to maintain the viable cell concentration within the optimal range (2×10^5^ - 5×10^5^ cells/ml). The cells were then split by dilution according to cell counts every 3-5 days until reaching the desired cell number. Cells were counted using a Vi-Cell Blu cell counter (Beckman Coulter) with trypan blue viability assessment.

On harvest day, cells from each line were pooled into a 250 ml conical tube, thoroughly mixed, and an aliquot of ∼10 ml was transferred to a T25 flask. An aliquot was removed to perform a cell count, and a sterility test was performed by streaking two sheep’s blood agar plates, then incubating the plates for two weeks at 30° C and 37° C. Based on the cell count, we determined the volume of cell suspension required to reach the desired cell number. These calculated volumes were transferred from each individual pool to a secondary tube for combining all cell lines. The pooled suspension was centrifuged at 228 RCF for 10 minutes, resuspended in 800 ml of cryopreservation medium (65% RPMI-1640, 30% FBS, and 5% DMSO), and dispensed into cryovials at a concentration of 5×10^6^ cells/vial. Cells were then cryopreserved using a controlled-rate freezer and stored in liquid nitrogen vapor.

#### Sequencing methods for the HapMap mixture

At the WashU-VAI Genome Characterization Center, the HapMap cell-line mixture underwent the following sequencing: 500× short-read WGS using Illumina NovaSeq X Plus and 100× long-read WGS using PacBio Revio.

### METHODS DETAILS

#### Short-read sequencing

Genomic DNA samples were quantified using the Qubit Fluorometer. Genomic DNA (∼600-1,000ng) was fragmented on the Covaris LE220 instrument targeting ∼375bp inserts.

Fragmented DNA was size selected using 0.8X ratio of Ampure XP beads (Beckman Coulter) to remove fragments less than 300bp. Dual indexed libraries were constructed utilizing the KAPA Hyper PCR-free Library Preparation kit (Roche Diagnostics, Cat # 7962371001). Full length custom adaptors were used during ligation (IDT, UDI/UMI configuration with 10bp UDIs and a 9bp UMI in the i7 position). Libraries were run with KAPA Library Quantification kit (Roche Diagnostics) to measure molar concentration. Libraries were sequenced on NovaSeq X using paired end reads extending 150bp. To achieve >500× coverage for this application, four libraries were constructed and sequenced with resulting data pooled from a mixture of the 4 library sources.

The molarity of each library was precisely determined through qPCR using the KAPA Library Quantification Kit according to the manufacturer’s protocol (KAPA

Biosystems/Roche) to generate appropriate cluster density for the Illumina NovaSeq X Plus instrument. Normalized libraries were sequenced on a NovaSeq X Plus Flow Cell using the 151×10×10×151 sequencing recipe according to manufacturer protocol, yielding >500× WGS coverage.

#### Long-read sequencing

PacBio HiFi SMRTbell libraries was prepared following PacBio protocol ‘Procedure & Checklist – Preparing Whole Genome and Metagenome Libraries Using SMRTbell Prep Kit 3.0’. Genomic DNA was fragmented with a mode of ∼20 kb using the Diagenode Megaruptor 3 instrument. Genomic DNA was initially processed with DNAFluid+ (P/N E07020001) using speed of 40 to dissociate aggregates and homogenize the DNA. The homogenized DNA was then sheared twice using Shearing kit (P/N E07010003) with speeds of 28 and 30.

Sheared sample was assessed via fluorometry (Qubit High Senstivity DNA Kit) and Agilent Femto Pulse (Genomic DNA 165kb Kit).

Libraries were made according to PacBio protocol utilizing barcoded adapters from the SMRTbell barcoded adapter plate 3.0 (PacBio P/N 102-009-200) to allow for multiplexing of samples during sequencing. Libraries were size selected using Sage PippinHT instrument and the 0.75% Agarose High-Pass 75E kit (P/N HPE7510) with a start size of 15,000 bp-17,000 bp. Size selected libraries were prepared for sequencing following instructions generated in PacBio SMRT Link v13.0 Sample Setup and utilizing PacBio Revio polymerase kit (P/N 102-817-600). Sequencing was performed on PacBio Revio sequencer using 30-hour movies with an ‘On Plate Concentration’ of 170pM-200pM. 3 SMRTcells were generated for a total of ∼100X/SMRTcell and 300Gb total coverage. Average Q-score values of Q33, Q33, and Q32 for the 3 SMRTcells were achieved.

#### Data processing

Illumina-generated short reads were preprocessed using fastp (v.0.23.2) with the parameters “--trim_poly_g --length_required 151” to filter polyG artifact reads. The processed reads were then aligned to Genome Reference Consortium Human Build 38 (GRCh38, GCA_000001405.15_GRCh38_no_alt_analysis_set.fna.gz) and Telomere-to-Telomere assembly of the CHM13 cell line (T2T-CHM13v2.0, GCA_009914755.4.fa.gz) using the BWA-MEM (v.0.7.17).^52,53^ Duplicated reads were marked using Picard (v.2.9.0) MarkDuplicates, followed by local realignment around indels position using GATK pipeline’s “RealignerTargetCreator” and “IndelRealigner” (v.3.8-1-0-gf15c1c3ef). Base quality scores were recalibrated using “BaseRecalibrator” and “ApplyBQSR” algorithms in GATK (v.4.1.0.0).^53^

PacBio HiFi long reads in unaligned BAM format were aligned to the same GRCh38 reference genome using pbmm2 (v.13.1) with “--sort”, “--strip” and “--unmapped” options. For the SV caller nanomonSV^54^, the unaligned raw BAMs were converted to FASTQ format using bam2fastq (v.3.1.1) and separately aligned to both GRCh38 and CHM13 reference genomes using minimap2 (v2.28-r1209) with the “-ax map-hifi” option. The resulting SAM files were subsequently converted to BAM format, sorted, and indexed using samtools (v1.9).

#### Benchmarking set generation

We generated a contig-level genome graph using the MC graph construction method (v2.9.7)^20^ with a reference assembly (either GRCh38 or T2T-CHM13) and twelve assemblies from six Hapmap samples in HG005, HG00438, HG002, HG02257, HG02486, and HG02622. HG005. The MC graph was deconstructed to VCF files in GRCh38 or CHM13 coordinates, vcfbub^19^ was then used to simplify variant representations. This was then used as the input of the benchmarking set. As a consequence of the graph constructing process, any overlapping variants were represented as multiallelic multi-nucleotide variants, so we first used “bcftools norm -m” (v1.21)^55^ to split these sites, and “rtg vcfdecompose --break-indels --break-mnps” (v3.12)^56^ to break apart the multi-nucleotide variants. To distinguish them from the initial gaps in the graph, which labeled as “.”, we labeled genotypes of the cell lines with overlapping other allelic variants in “*”. More detailed information was listed in the VCF headers.

With this processed VCF file, we combined the maternal and paternal haplotypes for each variant across all six samples. The combined VCF was then processed to categorize variants into three classes based on size and structure: SNVs (reference and alternate alleles = 1bp), indels (absolute length difference < 50bp), and SVs (absolute length difference between reference and alternate alleles ≥ 50bp). We filtered these separated variant files to only those within assembly-reliable regions. For T2T-CHM13, these assembly-reliable regions were regions flagged as haploid obtained from Liao et al. in all of the 12 assemblies^19^, while for GRCh38, we used “liftOver” function in R package “rtracklayer”^57^ with hs-to-hg38 chain file (hs1ToHg38.over.chain, UCSC Genome Browser) to liftover the reliable regions. Duplicated regions in GRCh38 caused by liftover were merged by “bedtools merge -d 1”.

To construct a somatic variant set from the HapMap mixture, we selected variants that were homozygous reference in germline variant source, HG005, and heterozygous or homozygous alternate in any of the other five somatic source samples. For sex chromosomes (X and Y), since HG005 was male with only one X and one Y chromosome, we included variants with hemizygous reference patterns (‘0|.‘ or ‘.|0‘), indicating a reference allele from either maternal or paternal haplotypes and unknown from the other. The variants were reformatted with appropriate headers, with genotype and expected VAF information added to the INFO field. Expected VAFs were calculated based on the mixture ratios; for example, a homozygous variant from HG02622 (10% in the HapMap mixture) that is absent in other cell lines would have an expected VAF of 0.1, while a heterozygous variant would have an expected VAF of 0.05. The final benchmarking set VCF files for SNVs, indels, and SVs were indexed and made available as separate resources https://wangcluster.wustl.edu/~juanfmacias/Graph_based_HapMap_Truth_Set/.

We built the MEI benchmarking set from the SV benchmarking set (**Figure S14**). Insertion-type SVs were extracted from the benchmarking set. Inserted sequences were annotated using RepeatMasker (v4.1.7; NCBI/RMBLAST 2.14.1+, Dfam 3.8)^58^, identifying MEIs classified as Alu, L1, and SVA elements. We further refined our MEI benchmarking set by selecting insertions belonging to recently active (“young”) mobile element subfamilies, specifically L1HS, L1PA2, L1P1, AluY subfamilies (AluY, AluYa5, AluYa8, AluYb8, AluYb9, AluYc, AluYd8, AluYf4, AluYg6, AluYh9, AluYk11, AluYk12, and AluYk4), SVA_F, SVA_E, and SVA_D.^22^ Lastly, inserted sequences were trimmed (MEI clipping) to retain only the MEI segment, resulting in the final MEI benchmarking set. In a few rare cases, an insertion SV contained more than one young MEIs. MEIs with the same chromosome and the same start position but different alternative allele sequences were listed in separate rows in the VCF file.

#### Additional Benchmark Resources

In addition to the nuclear genome, we include a mitochondrial benchmarking set encompassing SNVs and indels within chrM. To accommodate mitochondrial-specific characteristics, we did not apply HG005 variant filtering or reliable region masking. In this context, VAF is replaced by heteroplasmy level (HL), which was calculated based on the known mixture ratios and the presence of variants in maternal assemblies.

#### Sequencing simulation

Short-read sequencing data used for SNV and indel validation was generated using wgsim (0.3.1-r13).^59^ Wgsim was run with 151 read lengths, 345 insert size, 55 standard deviation, 0% mutation rate, 0% base error rate, 0% indel rate, and 0% indel extension rate settings. To generate HapMap mixtures with the desired coverage, we merged simulated FASTQ files from maternal and paternal assemblies for each HapMap sample, combining them according to predefined proportions (**Figure S15**).

Simulated short reads were aligned and processed following the same pipeline used for real HapMap Illumina sequencing data, as outlined in previous sections. The quality of the simulated data was assessed by samtools (v.1.10) “depth” and “stats” (**Table S1**).

Simulated long-read sequencing data was generated through two key steps: subread simulation and consensus sequence generation (**Figure S15**). First, pbsim3^60^ was utilized with a WGS strategy. The “qshmm” method was employed as the error model, using the provided QSHMM-RSII.model. For each library, sequencing depth was calculated based on a predetermined mixture ratio, with the “pass-num = 7”. Fragment length parameters were defined as follows: minimum 300 bp, maximum 80,000 bp, mean 22,000 bp, and standard deviation 5,000 bp. To simulate errors, all introduced sequence errors were set as substitutions using the option “--difference-ratio 1000:0:0”. Next, subreads were processed into consensus sequences using PacBio’s software tools: pbindex v3.1.1 and ccs v6.4.0, both run with default parameters.

#### Alignment-based validation

We created two pipelines to validate the presence of variants in our benchmarking set in the simulated data (**Figure S16**).

SNVs: mpileup+ bcftools isec

We used bcftools mpileup to obtain counts of every SNV present in a simulated BAM file using bcftools v 1.10.2.^61^ mpileup was run with the paramaters “-a FORMAT/AD,FORMAT/DP, -d 20000 -q 0 and -Q 0”

To decrease the runtime of mpileup, we subsetted the benchmarking set by chromosome and ran mpileup on each chromosome in parallel, using bcftools (v1.10.2) “merge” to join the subsetted output VCFs. Following the run, we subsetted the VCF to only the SNVs called and used bcftools (v1.10.2) “isec” to compare the mpileup VCF to the benchmarking set.

Indels and SVs: cigar+travri

For both indels and SVs, we created a python script to extract all the insertions and deletions from the cigar codes of the aligned BAM files. Briefly, we subsetted the simulated BAM file into reads that overlap with a called indel or SV in the benchmarking set, plus or minus 50 bases using “samtools (v1.10) view”. For each subsetted BAM file, we identified where insertions and deletions were using CIGAR codes, and then used a reference FASTA file, the position of a read, and sequence of a read to create a list of all the SVs and indels present. We then used “bcftools (v1.10.2) merge” to merge all these variant lists from different regions into a single list of all indels/SVs present. Given that a read needs to span the entire variant to detect it, we used simulated short-read sequencing data for indel detection and simulated long-read sequencing data for SV detection. We used Truvari^24^ (v4.0.0) with “–s 50 –S 50 – pctseq 0.7 –pctsize 0.7” for SVs, and “–s 1 –S 1 –pctseq 1 pctsize 1” for indels to compare this to our benchmarking set, and record the output precision.

### Assembly-based validation

We used Dipcall^25^ to perform pairwise comparisons between the haplotype-resolved assemblies from the six HapMap individuals to identify SNVs, indels, and SVs. Here, HG005 was designated as the germline reference because it constitutes 83.5% of the HapMap mixture. For the remaining five samples, we compared each of their 10 assemblies (maternal and paternal) to the HG005 maternal and paternal assemblies respectively, resulting in 20 VCF files containing all types of variants from the comparisons.

Because Dipcall performs pairwise alignment through minimap2^62^ whereas the graph-based Cactus alignments are conditioned on all prior alignments, the same variant can be represented differently by the two methods (**Figure S8b**). To reconcile the differences, we used an in-house python script to reconstruct each sample’s sequence around the variant from both the graph and Dipcall output using reference genome sequence. We considered a variant validated if the somatic variant in the graph appears in the Dipcall variant list or if the two reconstructed sequences are the same.

### Distortion matrix

To contextualize deviations between expected and observed VAFs, we apply a distortion mapping framework to quantify how alignment to a reference genome perturbs the placement of reads. We simulate 150 bp paired-end reads from haploid query assemblies using wgsim^59^, and align them to the target reference genome using BWA-MEM^52^. For each aligned read, we record the query interval from which it was simulated (the source interval) and the interval in the reference where it aligns (the target interval). To enable comparison in a shared coordinate system, we align each haploid assembly to the reference using minimap2^62^ and use paftools to lift source intervals into reference coordinates. This process is repeated independently for each haplotype (maternal and paternal) of each individual in the mixture.

To capture regional patterns, we define 50 kb sliding intervals along the reference and aggregate all simulated reads whose lifted source and aligned target positions fall within those intervals. For every interval, we count how many reads originating from each lifted source interval align to each target interval, producing a source–target matrix. Each column of this matrix is normalized to sum to one, yielding a conditional probability distribution over target intervals for a given source interval. The diagonal of this normalized matrix reflects the fraction of reads that remain within their original interval after alignment. From this, we compute two distortion metrics per interval: diagonal distortion, defined as one minus the fraction of reads that remain in the same interval after alignment, and coverage distortion, defined as one minus the total read placement probability assigned to the interval, including both reads that originated from the interval and remained there and reads that mapped there from elsewhere. An interval is labeled distorted if either metric exceeds 0.05.

This approach produces haplotype- and interval-resolved distortion maps that highlight

reference-induced perturbations in read placement. While the current study focuses on reference-based alignment, these maps provide a framework for identifying intervals where observed VAF deviations may be attributable to technical artifacts introduced by the reference, rather than true underlying biological variation.

### Modeling somatic variant detection rates

We built a suite of six stochastic models to predict probability of detecting a somatic variant in at least *k_0_* reads out of *N* total reads from the variant’s underlying allele frequency.

Throughout, we assume simple random sampling without replacement and each read is an independent draw whose success probability noted as _obs_

Ideal sampling (no explicit error)

- Model 1 – Binomial

If sequencing is error-free, the number of reads with alternative allele (*K)* follows a binomial distribution.

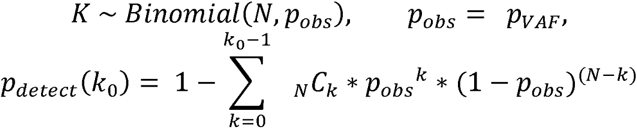

Binomial error extensions

Sequencing introduces substitution errors with per-base rate *p_error_*. A reference read can be mis-called as the mutant allele (false positive), and a mutant read can revert to reference (false negative). Assuming an equal probability of the three incorrect nucleotides, the effective success probability per read becomes

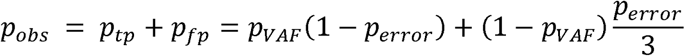

- Model 2 – Binomial +error

Replace *p_obs_*in the binomial formula above.

Poisson error extensions

Empirically, false-positive mutant reads occur independently across genomic sites, closely matching a Poisson process. Let λ be the expected number of erroneous mutant reads per site,

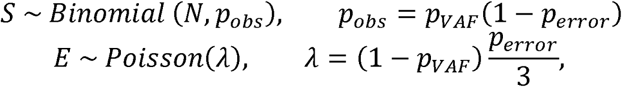

- Model 3 – Binomial sampling + Poisson error.

Total mutant calls *K = S+E*. Because *S* and *E* are independent,

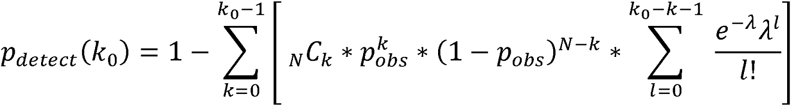

Using the HapMap-mixture dataset generated from other SMaHT Genome Characterization Centers downloaded from SMaHT data portal, we estimated empirical detection rates across the full VAF spectrum and computed the root-mean-square error (RMSE) between each

model’s predictions and observation (**Figure S10**). To identify which model best explains the observed data, we compared the RMSE between each model’s predictions and the empirical data (**Figure S10c**). The model yielding the lowest RMSE was selected as the best fit. We then used this best-fit model to generate a heatmap illustrating the number of sequencing coverage required to achieve specific detection rates at various VAFs (**Figure S11**). All formulas were implemented in MATLAB R2023a.

### Assessing the performance of somatic variant callers

We called somatic SNVs and indels using Strelka2 (v2.9.7)^29^, Varscan2 (v2.4.2)^30^, and Mutect2 (GATK v4.4.0.0)^27^ in GRCh38 and CHM13 aligned 500× WashU-sequenced short read data. Somatic SVs were called using Sniffles2 (v2.2)^28^ and Severus (v0.1.1)^31^ in 100× WashU-generated long-read sequencing data. Somatic MEIs were called using PALMER (v2.0.0)^32^ and xTEA long release (v0.1.0)^33^ in 100× WashU-generated long read sequencing data. Detailed commands can be found in **Table S4** To evaluate caller performance exclusively on somatic variants, we employed “tumor-normal” comparison mode for Strelka2,

Varscan2, and Mutect2 using 420× simulated short-read data generated with wgsim^59^ from the HG005 assemblies and for Severus using 100X simulated long-read data generated with pbsim3^60^. For callers lacking a tumor-normal mode, including Sniffles2, PALMER, and xTEA long-read mode, we conducted two separate runs with HapMap real data (as “tumor”) and HG005 simulated data (as “control”), subsequently selecting unique variants found exclusively in HapMap samples using Truvari^24^.

To compare our benchmarking set against variants identified by each caller, we used vcfeval option with “--squash-ploidy” of RTGTools^56^ for SNVs and indels, Truvari ^24^ (v4.0.0) with “–s 50 –S 50 –pctseq 0.7 –pctsize 0.7 --pick multi” for SV and Truvari ^24^ (v4.0.0) with “–s 50 –S 50 –pctseq 0.1 –pctsize 0.1 --refdist 50” for MEIs (**Figure 4**). We define the benchmark region as the assembly reliable region mentioned in the **Benchmark set generation** section. Variants identified by callers but not present in the benchmarking set were classified benchmarking false positives, variants present in both the truth set and callers were true positives, and truth set variants not identified by callers were false negatives.

Variants outside these defined regions were excluded from benchmarking assessments.

To evaluate caller performance across VAFs, we first binned the benchmarking set by VAF and assessed overlaps with each caller’s calls to calculate recall. For precision, we binned each caller’s detected variants by VAF and measured their overlap with the full tr benchmarking uth set. Caller performance was further compared with respect to genomic mappability using UMAP-derived K-mer metrics.^34^ Specifically, we assessed single-read mappability of GRCh38 and T2T-CHM13 reference genomes, defined as the fraction of a genomic region overlapping at least one uniquely mappable 24-mer, 36-mer, 50-mer, or 100- mer. Genomic regions were stratified into hierarchical, non-overlapping categories: uniquely mappable k24-mer regions; uniquely mappable k36-mer regions excluding overlaps with k24; uniquely mappable k50-mer regions excluding k24- and k36-mer regions; uniquely mappable k100-mer regions excluding k24-, k36-, and k50-regions; and remaining regions lacking unique mappability across these lengths.

### Assessing the performance of somatic variant callers

To analyze regional biases in variant calling performance, we partitioned the human genome into non-overlapping 1-kb windows and classified each window based on variant content. SV harboring regions were defined as windows containing at least one benchmark SV, while regions with no true positive calls (noTP regions) were defined as windows where variant callers failed to detect any benchmark variants correctly despite their presence.

Challenging genomic features were obtained from multiple publicly available resources. Last 5 Mbp of each chromosome was calculated based on chromosome lengths defined in NCBI Genome assembly GRCh38.p14 (https://www.ncbi.nlm.nih.gov/datasets/genome/GCF_000001405.40/). Centromere regions were obtained from NCBI Genome Reference Consortium (https://www.ncbi.nlm.nih.gov/grc/human). All other regions were obtained from Genome-in-a-bottle genome stratifications v3.5 (https://ftp-trace.ncbi.nlm.nih.gov/ReferenceSamples/giab/release/genome-stratifications/v3.5/GRCh38@all/). Specifically, in **Figure 5 and S13**, we used GRCh38_AllTandemRepeats.bed.gz for “TandemRepeats”, a concatenation of simple repeats homopolymer, di-, tri-, and quad-TR files for “SimpleRepeats”, GRCh38_satellites_slop5.bed.gz for “Satellites”, GRCh38_gclt25orgt65_slop50.bed.gz (GC content less than 25% or greater than 65%) for “AbnormalGC”, and GRCh38_segdups_gt10kb.bed.gz (segmental duplications with length greater than 10 kb) for “SegDup”.

For each genomic region category, we calculated the proportion of 1-kb windows overlapping with specific challenging features across three sets: the entire genome, SV-harboring regions, and noTP regions. Fold change enrichment was computed as the ratio of proportions between comparison sets. To account for potential coverage limitations affecting low-VAF variants, we performed a secondary analysis restricting SV-harboring regions to those containing benchmark variants with ≥10% variant allele frequency, with noTP regions redefined accordingly within this subset.

### Coverage calculation for clinical relevant genes

For the special case where the detection threshold is a single supporting read (), the general binomial model with error-adjusted success probability simplifies to a closed exponential form. Specifically, the probability of nondetection corresponds to the event of observing zero alternate reads,

The detection probability is then

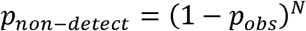

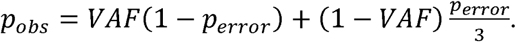

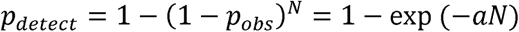

For and _error_ ^-6^ this evaluates to _obs_ and . Thus, under a single-read threshold, detection probability follows a one-parameter exponential rise,

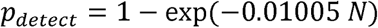

which provides an analytically convenient form for describing coverage–detection relationships.

To capture heterogeneity across the genome, we fitted this exponential model separately for each gene using benchmark SNVs at 1% VAF. This per-gene fitting allowed us to account for local sequence contexts and coverage variability that influence effective detection rates. In practice, we re-parameterized the exponential model as

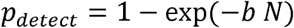

where b serves as a gene-specific weighting factor that encodes the combined influence of VAF, sequencing error, and local genomic features on detection probability. By estimating b from observed detection rates in each gene, we obtained gene-level coverage requirements for achieving specified sensitivity thresholds (e.g., 80% at 1% VAF). Observed detection rates were obtained from Mutect2 (tumor-only mode, default parameters) applied to Illumina NovaSeq X Plus sequencing data generated by four GCCs (WashU, BCM, NYGC, and Broad). These fitted parameters form the basis of our per-gene coverage estimates (Table S6).

## SUPPLEMENTARY FIGURES

**Figure S1.**
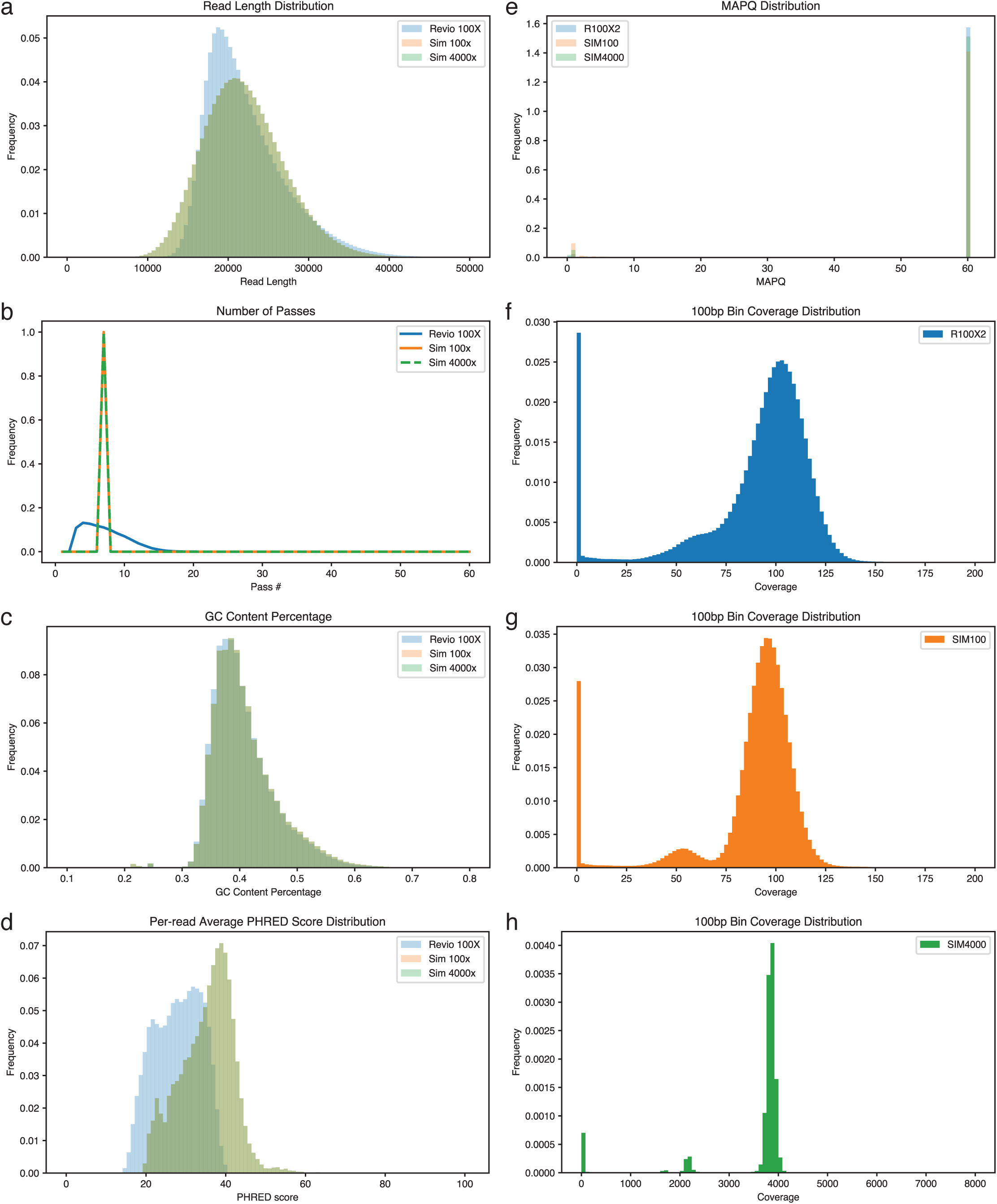
Quality control for HapMap long-read sequencing data. For real 100x long-read sequencing data, simulated 100x long-read data, and simulated 4000x long-read data, we plotted the distribution of their (a) read length, (b) number of passes, (c) GC content, (d) per-read base quality scores, (e) mapping quality, (f) coverage calculated per 100bp bins for real 100x long-read, (g) coverage calculated per 100bp bins for simulated 100x long-read, and (h) coverage calculated per 100bp bins for simulated 4000x long-read data.

**Figure S2.**
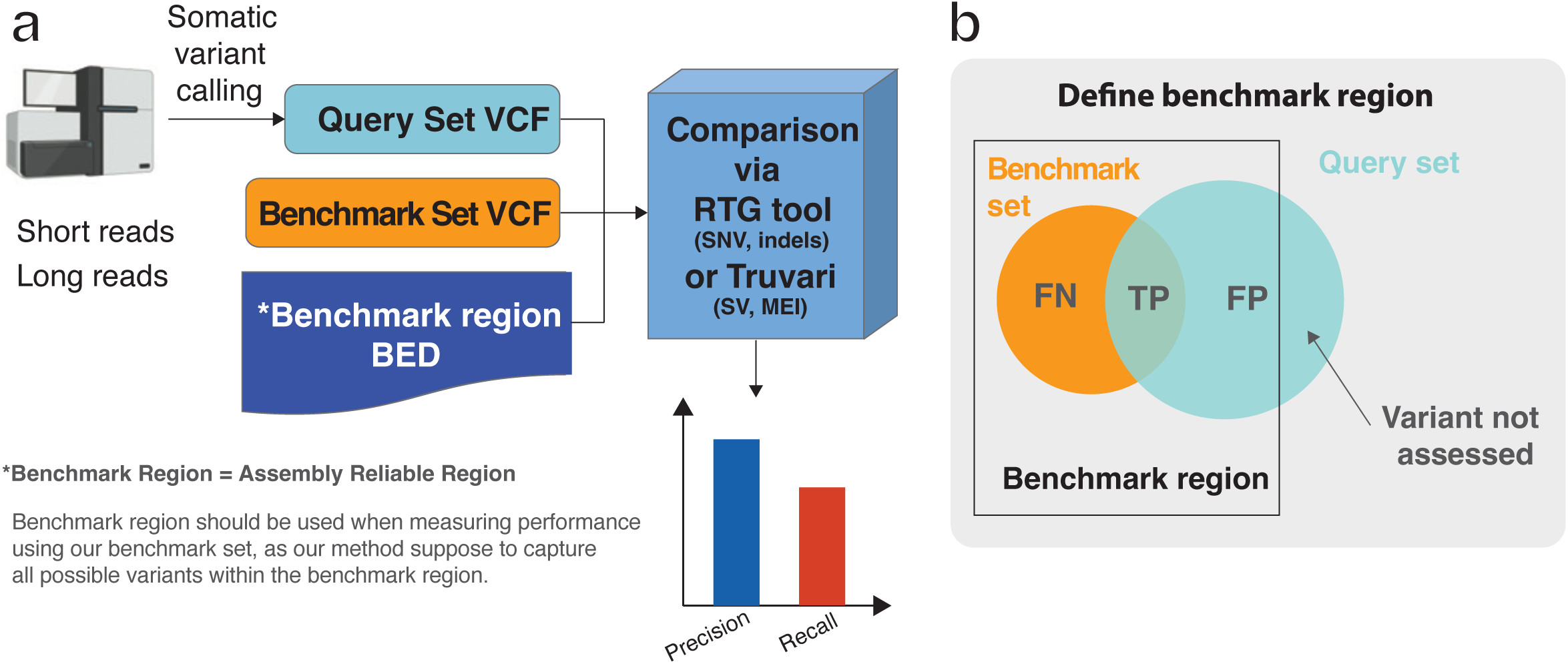
Definition of benchmarking regions and evaluation metrics. (a) Workflow for somatic variant calling evaluation using query set VCF, benchmarking set VCF, and benchmark region BED files. (b) Venn diagram illustrating the relationship between benchmarking set and query set variants within and outside the benchmark region, with true positives (TP), false negatives (FN), and false positives (FP) labeled correspondingly.

**Figure S3.**
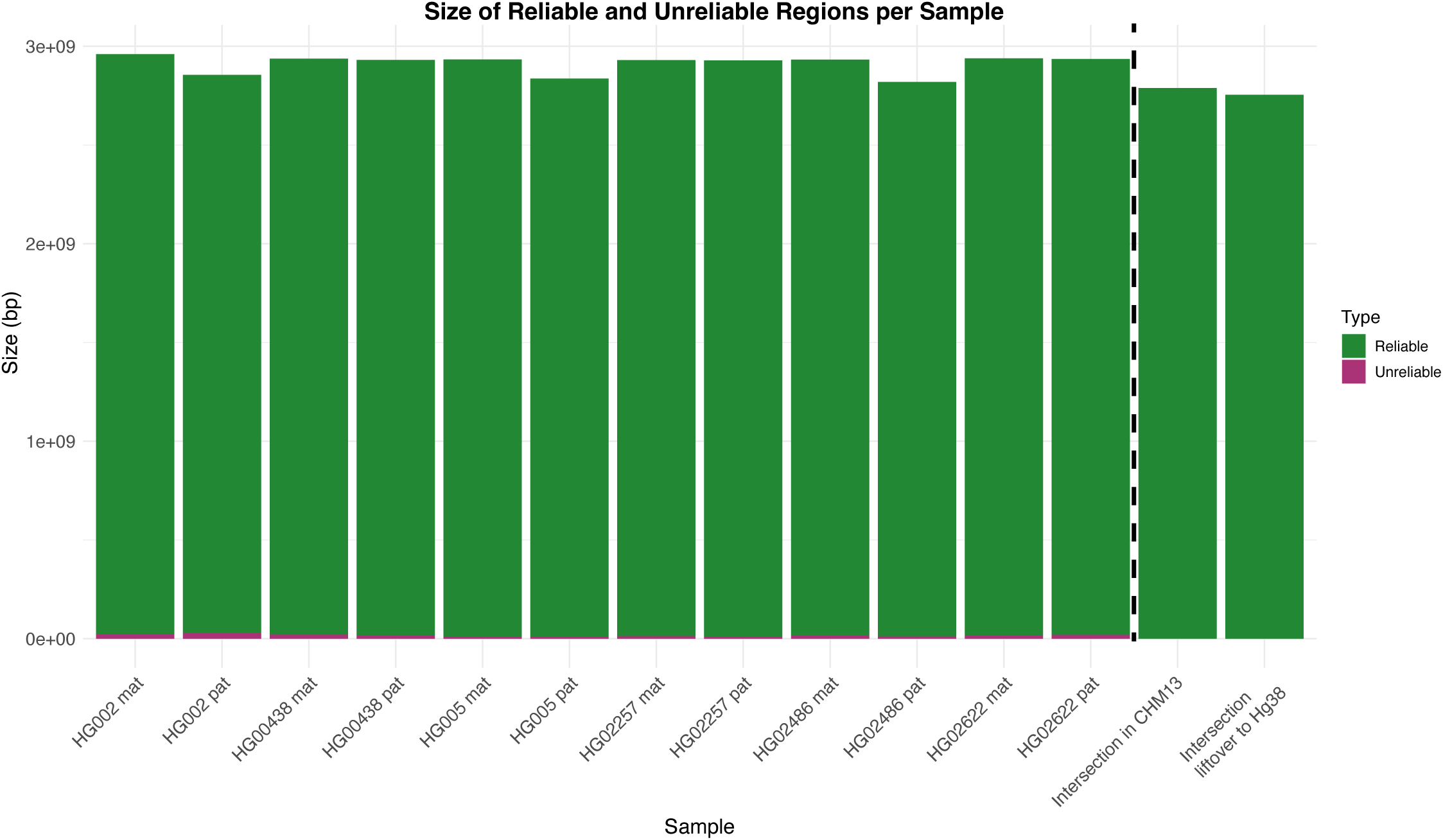
Characteristics of variants at the assembly level. Quantification of reliable (green) and unreliable (purple) genomic regions for each haplotype assembly, including both maternal (mat) and paternal (pat) haplotypes. The two rightmost columns show the consensus reliable regions across all assemblies in CHM13 and GRCh38 (after liftover) coordinates.

**Figure S4.**
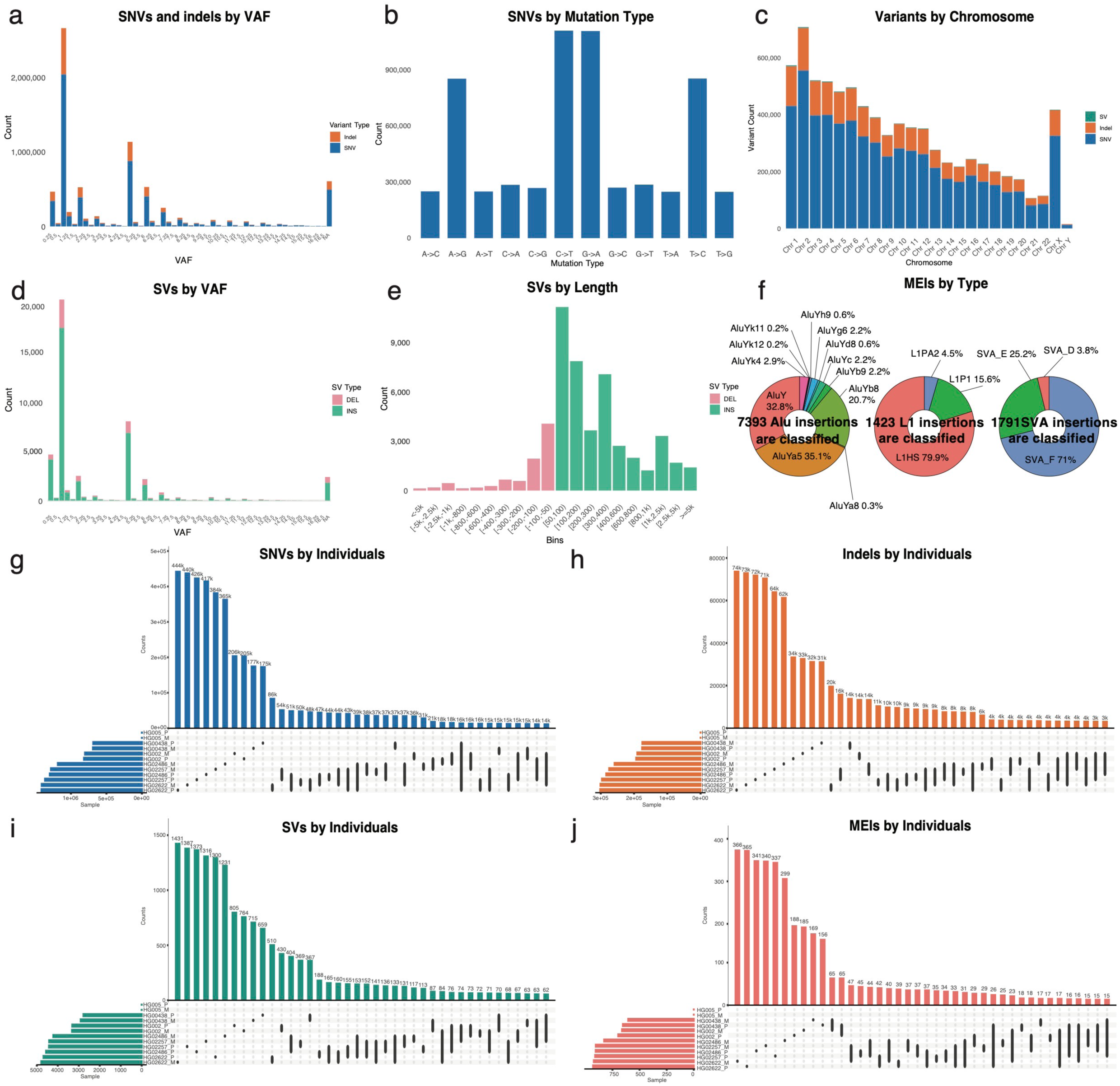
Characteristics of variants in the Hg38-based benchmarking set. (a) Number of SNVs (blue) and indels (orange) by expected VAF. (b) SNV counts categorized by mutation type. The benchmarking set contains more transitions than transversions. (c) Number of SNVs (blue), indels (orange), and SVs (green) across all autosomal and sex chromosomes. (d) Number of SVs, with insertion events in green and deletion events in pink, by different expected VAFs. (e) Number of SVs, with insertion events in green and deletion events in pink, by different lengths. (f) Distribution of different types of MEIs by subfamily. (g)-(j) Upset plot of number of SNVs (g), indels (h), SVs (i), and MEIs (j) per haplotype assembly of the HapMap individuals. VAF, variant allele frequency; SNV, single nucleotide variant; indels, small insertions and deletions; SV, structural variant; MEI, mobile element insertions.

**Figure S5.**
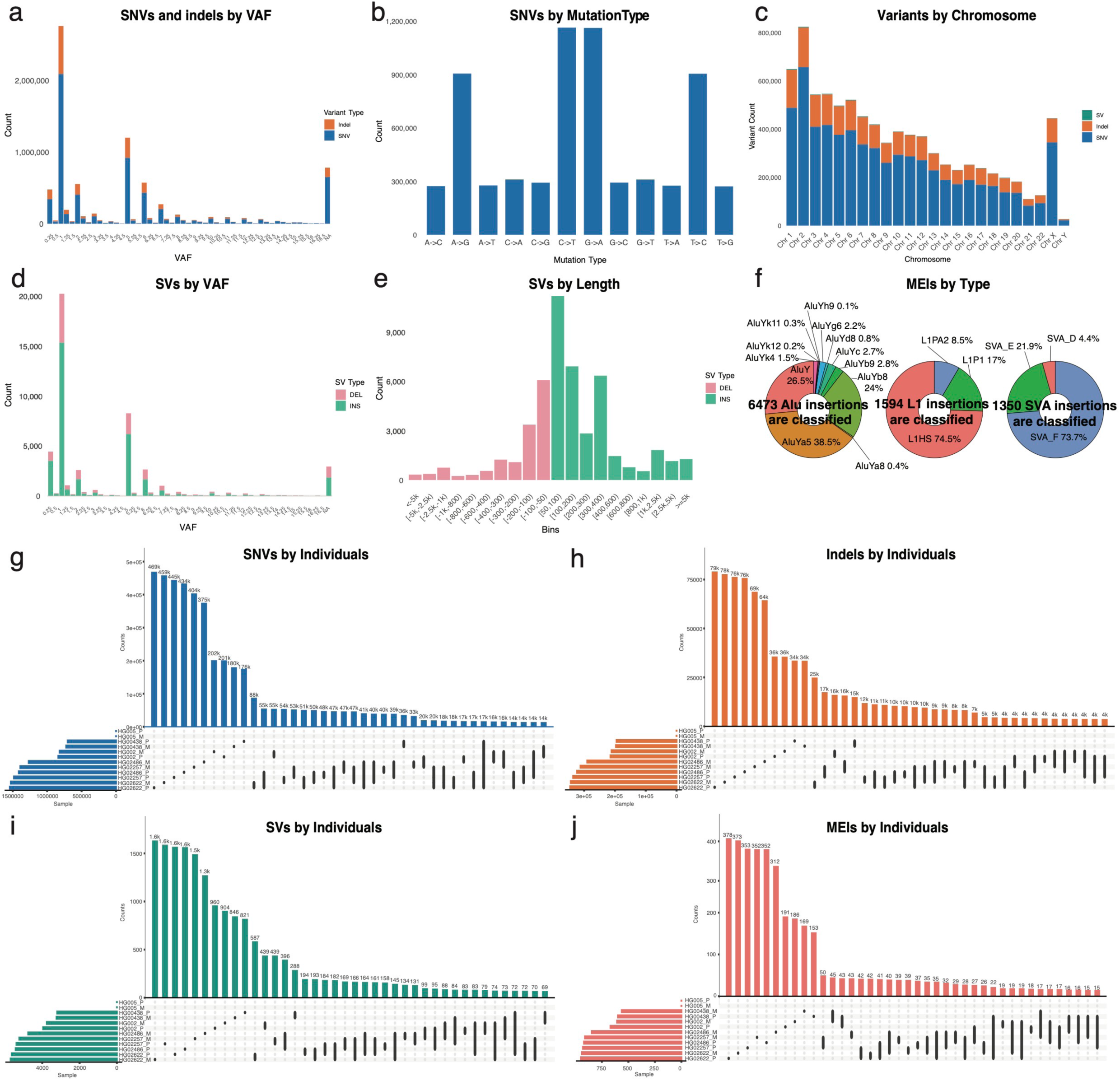
Characteristics of variants in the CHM13-based benchmarking set. (a) Number of SNVs (blue) and indels (orange) by expected VAF. (b) SNV counts categorized by mutation type. The benchmarking set contains more transitions than transversions. (c) Number of SNVs (blue), indels (orange), and SVs (green) across all autosomal and sex chromosomes. (d) Number of SVs, with insertion events in green and deletion events in pink, by different expected VAFs. (e) Number of SVs, with insertion events in green and deletion events in pink, by different lengths. (f) Distribution of different types of MEIs by subfamily. (g)-(j) Upset plot of number of SNVs (g), indels (h), SVs (i), and MEIs (j) per haplotype assembly of the HapMap individuals. VAF, variant allele frequency; SNV, single nucleotide variant; indels, small insertions and deletions; SV, structural variant; MEI, mobile element insertions.

**Figure S6.**
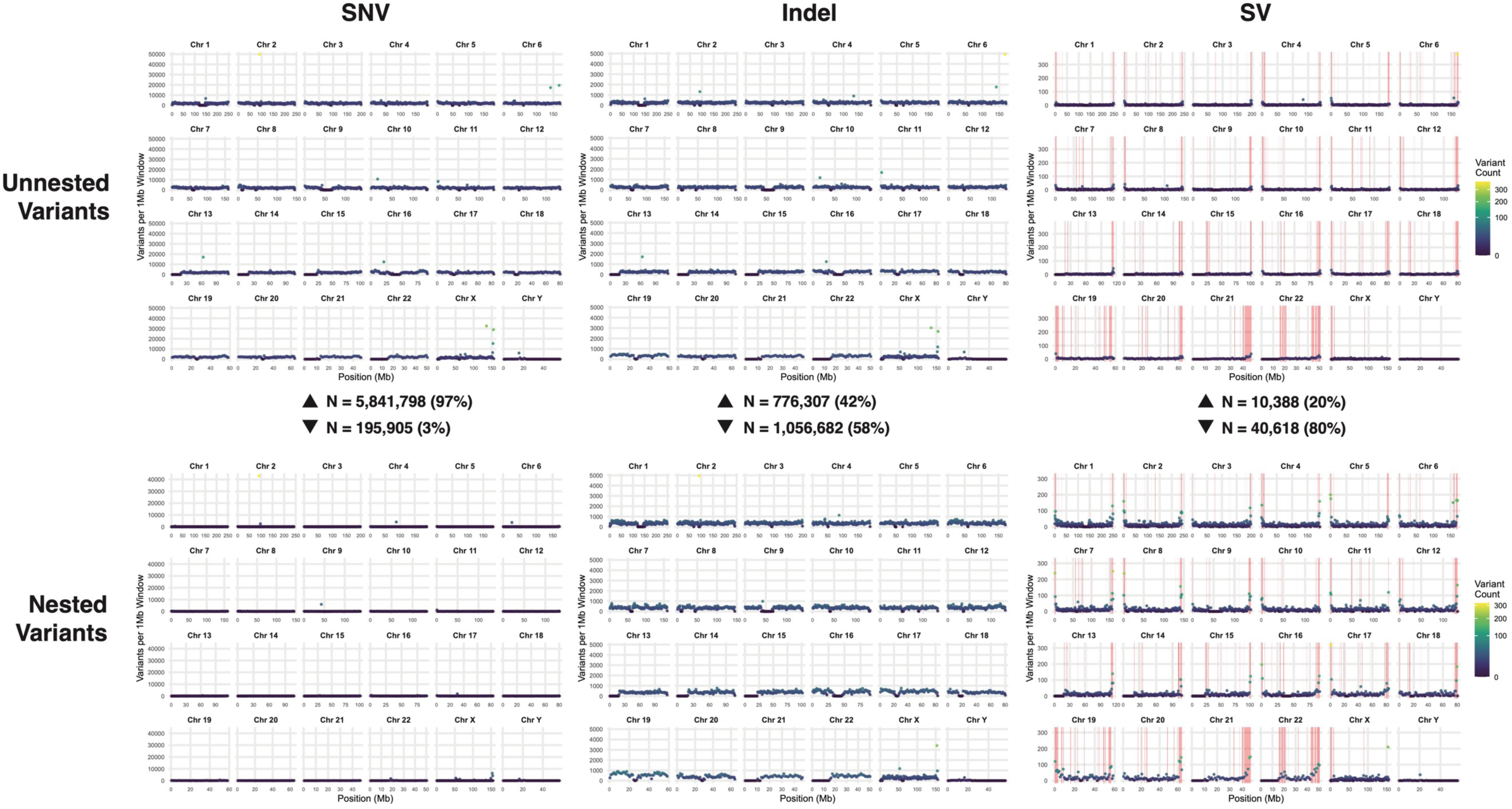
Mutation hotspot of SNV, indel, and SV in Hg38-based benchmarking set. We measured the number of nested (variants overlapping with others) and unnested (variants do not overlap with others) variants in each 1Mb window. 97% of SNV, 42% of indel, and 20% of SV are unnested. No variant hotspots were observed in SNV or indel, but hotspots were observed in SV and overlap largely with the SV hotspots reported by the Human Genome Structural Variation Consortium. SNV, single nucleotide variant; indels, small insertions and deletions; SV, structural variant.

**Figure S7.**
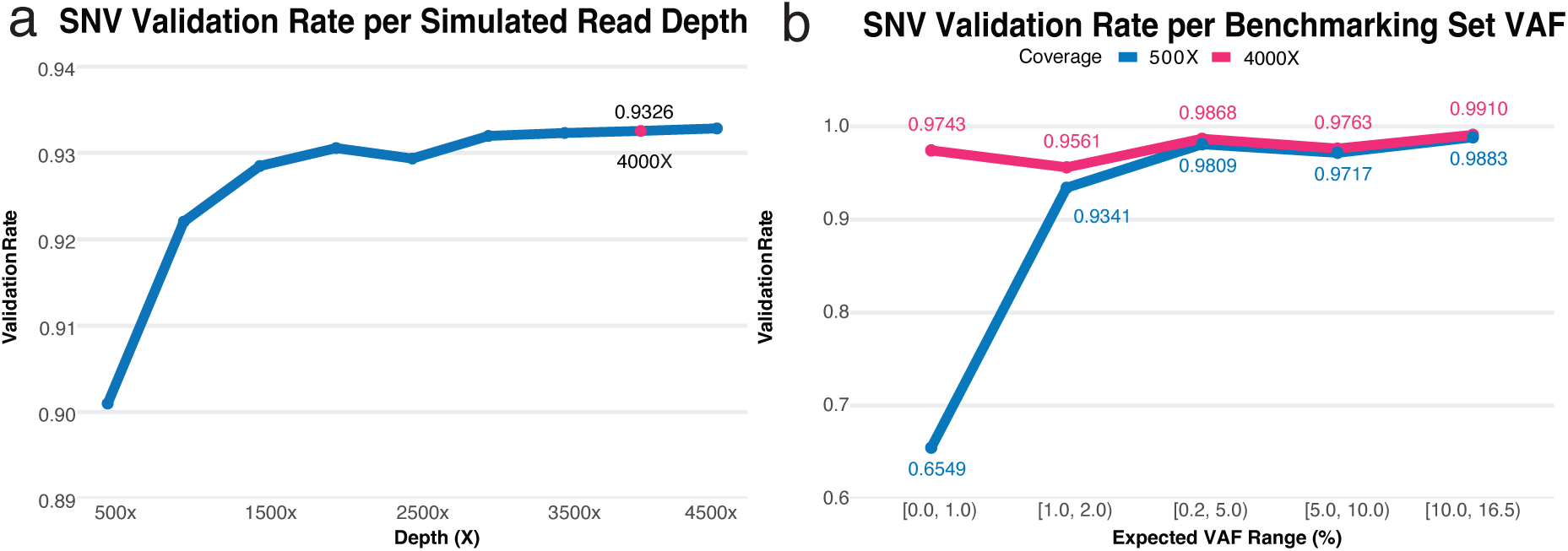
Validation of the somatic benchmarking set. (a) SNV validation rates across different simulated read depths from 500× to 4500×, with validation rate plateauing at around 4000×. (b) SNV validation rates across different VAF ranges. Validated from 4000× sequencing is in pink, and 500× sequencing is in blue. Validation rates drop for low VAF variants in 500× data but not 4000× data.

**Figure S8.**
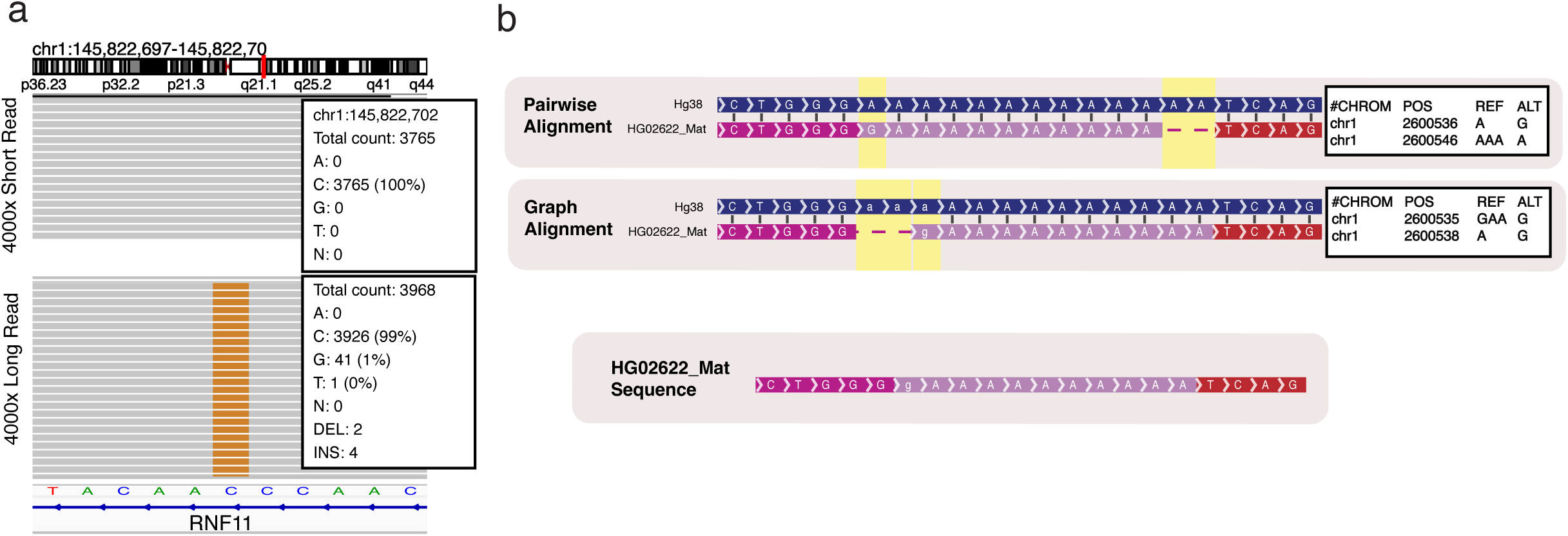
Example of true variant missed by alignment-based validation. (a) In the top track, short read sequencing does not identify any reads that have a G in the given position. However, in the bottom track, long read sequencing, which is less likely to be misaligned, shows the C to G mutation in 1% of the reads. (b) The top track represents the Dipcall pairwise alignment of HG02622_Mat to hg38. The two-base-pair deletion in places at the beginning of the A repeat, with the A to G mutation immediately following. However, in the bottom track, representing graph’s joint alignment of HG02622_Mat, the two-base-pair AA deletion is aligned to the end of the A repeat and the A to G mutation is aligned to the start. Both forms of alignment reflect the same sequence composition, and therefore both are valid alignments of the sequence and represent the same mutations.

**Figure S9.**
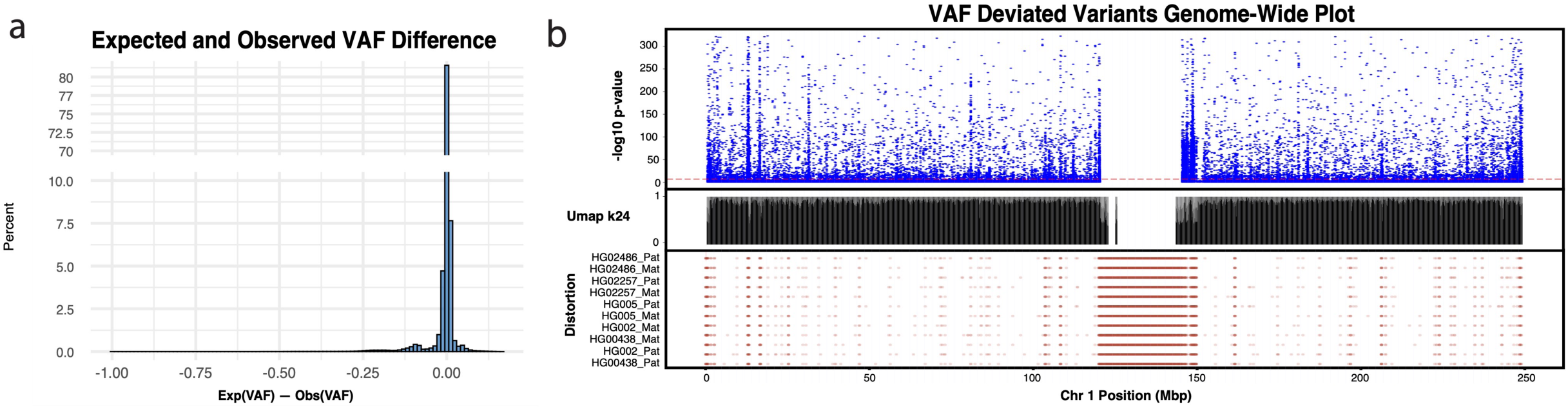
Validation of the somatic benchmarking set. (a) Distribution of differences between expected and observed VAFs for benchmarking set variants validated by the alignment-based method. (b) Top: Manhattan plot showing -log10 P-value of observed VAF deviated from expected VAF for all variants in chromosome 1; Middle: UMAP of 24-mers mappability across chromosome 1; Bottom: distortion metric distribution of all variants across chromosome 1. Variants reside in distortion regions exhibit significantly higher deviation between expected and observed VAF values (Pearson’s chi-squared test, p < 2.2 × 10LJ¹LJ). SNV, single-nucleotide variant; expVAF, expected variant allele frequency; obsVAF, observed variant allele frequency; UMAP, unique mappability.

**Figure S10.**
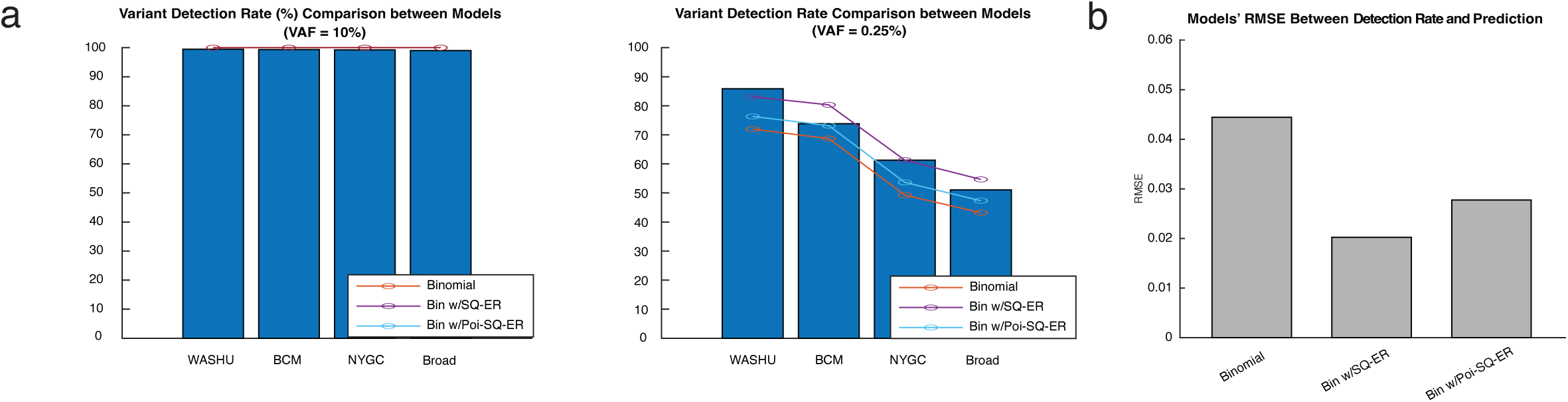
Model fitting for somatic variant detection rate in sequencing data from different sequencing centers. (a) Variant detection rate predictions from various models are similar for high VAF variants (VAF = 10%), and closely related to the real detection rate in the benchmarking set. (b) Variant detection rate predictions vary in between various models in low VAF variants (VAF = 2.5%). (c) Root mean square error (RMSE) calculated between the predictions made by the three models and the detection rate using the benchmarking set. Binomial distribution with sequencing error (Bin w/SQ-ER) shows the lowest RMSE. VAF, variant allele frequency.

**Figure S11.**
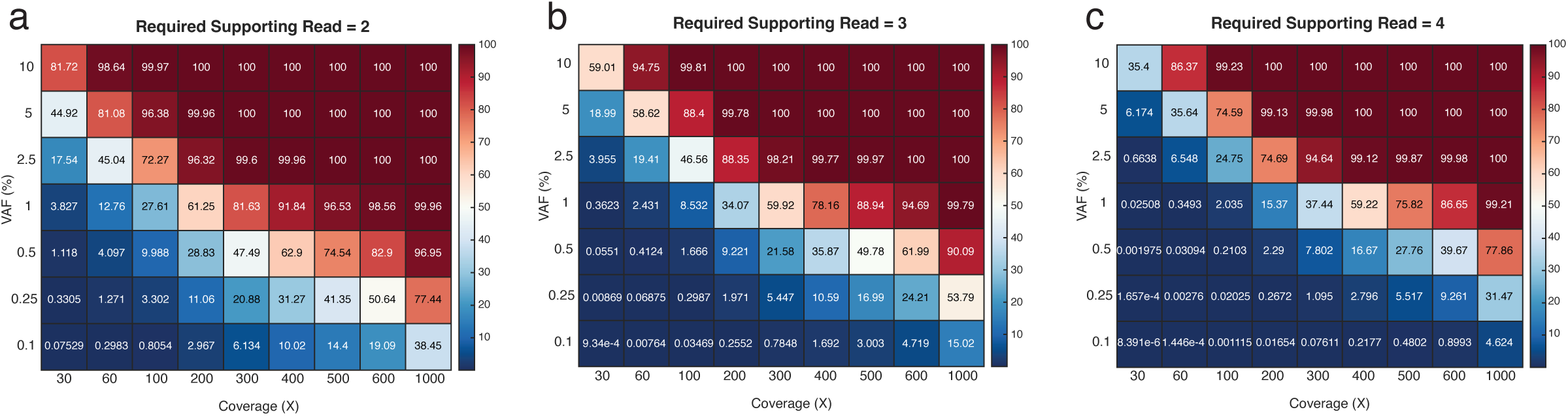
Optimal sequencing depth allowing minimum N reads with alt allele. Heatmaps of expected recall for different VAF and read depths when requiring one supporting read (k0=1), two reads (k0=2) three reads (k0=3) and four reads (k0=4). Boxes in red are closer to 100% recall while boxes in blue are closer to 0% recall.

**Figure S12.**
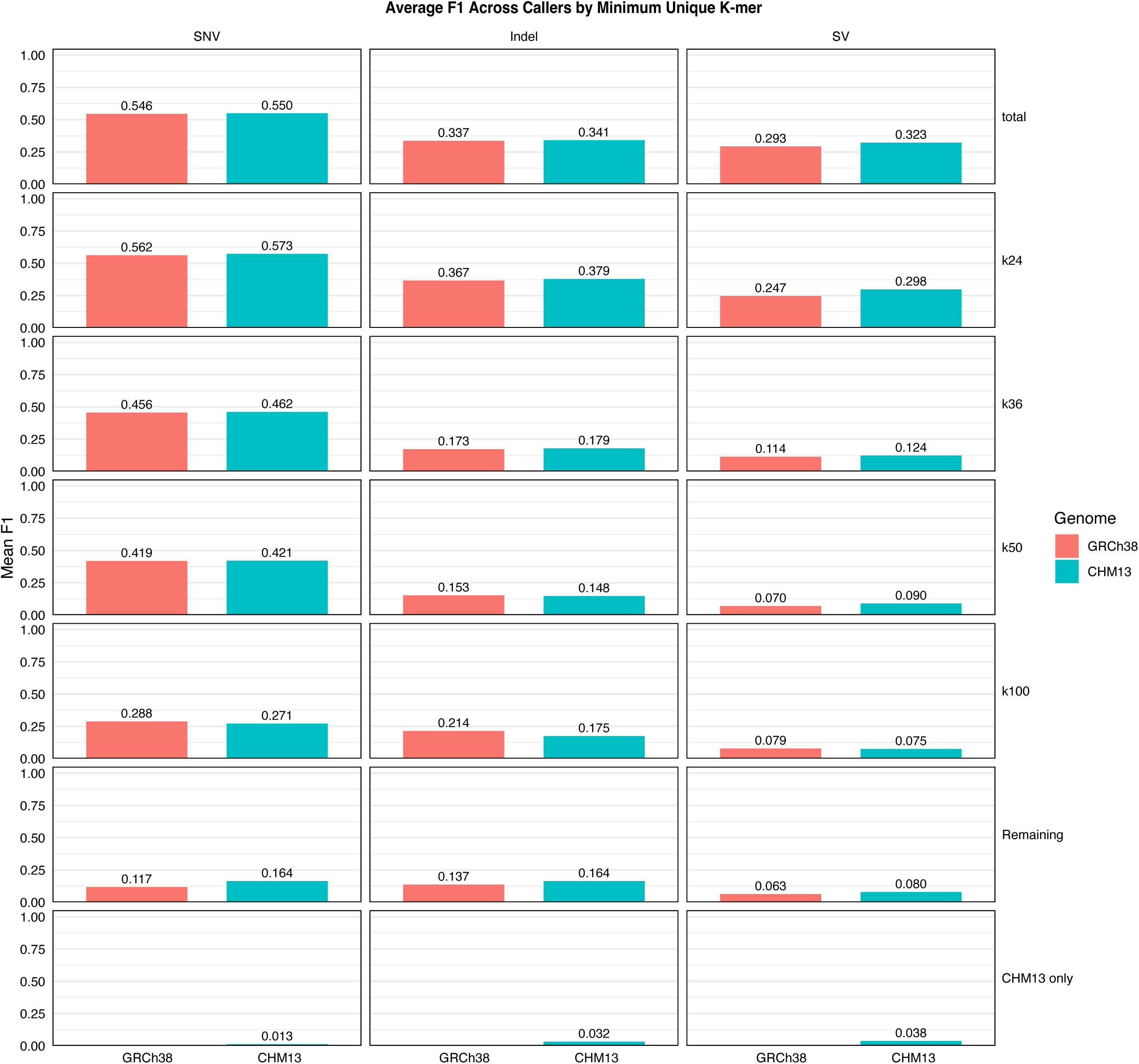
Variant caller performance across reference genomes stratified by genomic complexity. Average F1 scores across all variant callers for SNV, indel, and SV comparing GRCh38 and CHM13 reference genomes. Performance metrics are stratified by minimum unique k-mer length categories representing increasing genomic complexity: total (all regions), k24, k36, k50, k100, remaining (regions with k-mer lengths above k100), and CHM13-only (regions absent in GRCh38). SNV and indel detection showed comparable performance between reference genomes across all complexity categories. SV detection demonstrated reference-dependent performance differences, with CHM13 provided improved average F1 scores. Performance declined substantially in genomically complex regions (higher k-mer thresholds) for all variant types, with the most pronounced effects observed for SVs. SNV, single nucleotide variant; indels, small insertions and deletions; SV, structural variant.

**Figure S13.**
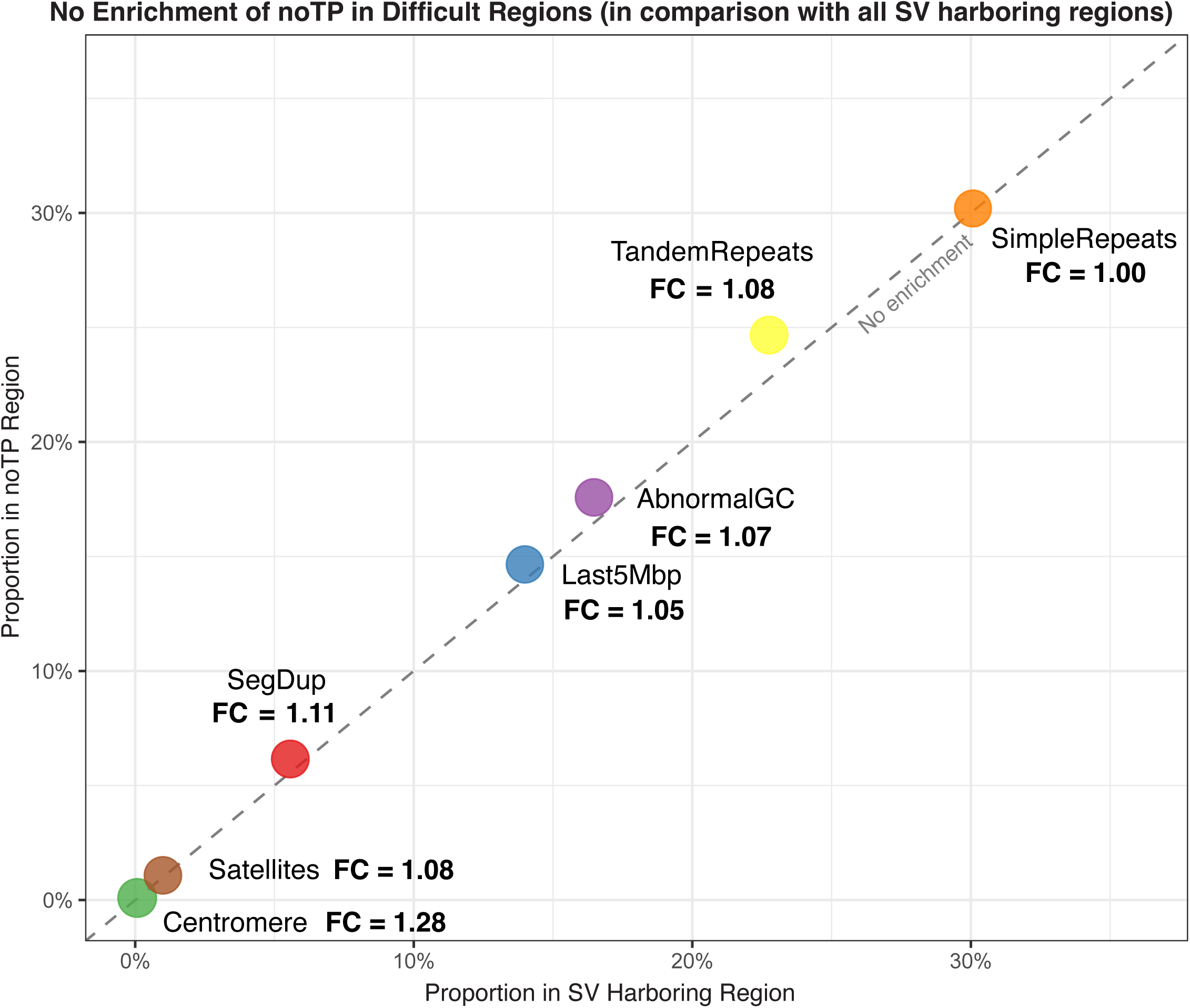
Absence of specific enrichment patterns for detection failures across challenging genomic regions. Enrichment analysis comparing noTP regions against all SV-harboring regions without VAF filtering. All challenging genomic features cluster near the diagonal line (FC ≈ 1.0), indicating no substantial enrichment of detection failures in any specific difficult sequence context. This suggests that variant caller failures occur broadly across various challenging regions rather than concentrating in particular sequence features. FC, fold change.

**Figure S14.**
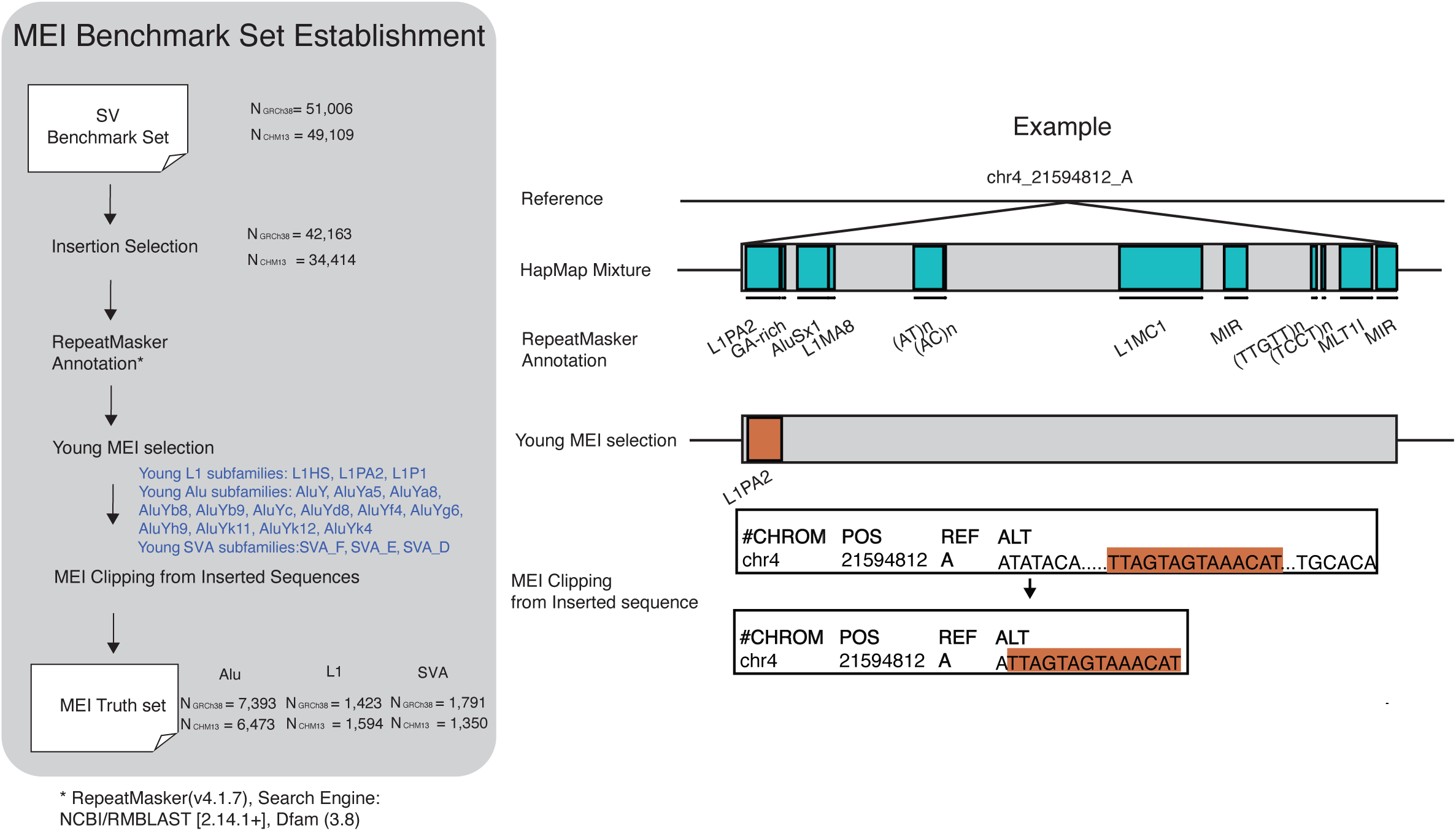
MEI benchmarking set generation. Process of establishing somatic MEI benchmarking set from SV set. Workflow is shown in the left panel. One example of different MEIs within the same inserted sequence at different annotation and filtering step is shown on the right. MEI, mobile element insertion.

**Figure S15.**
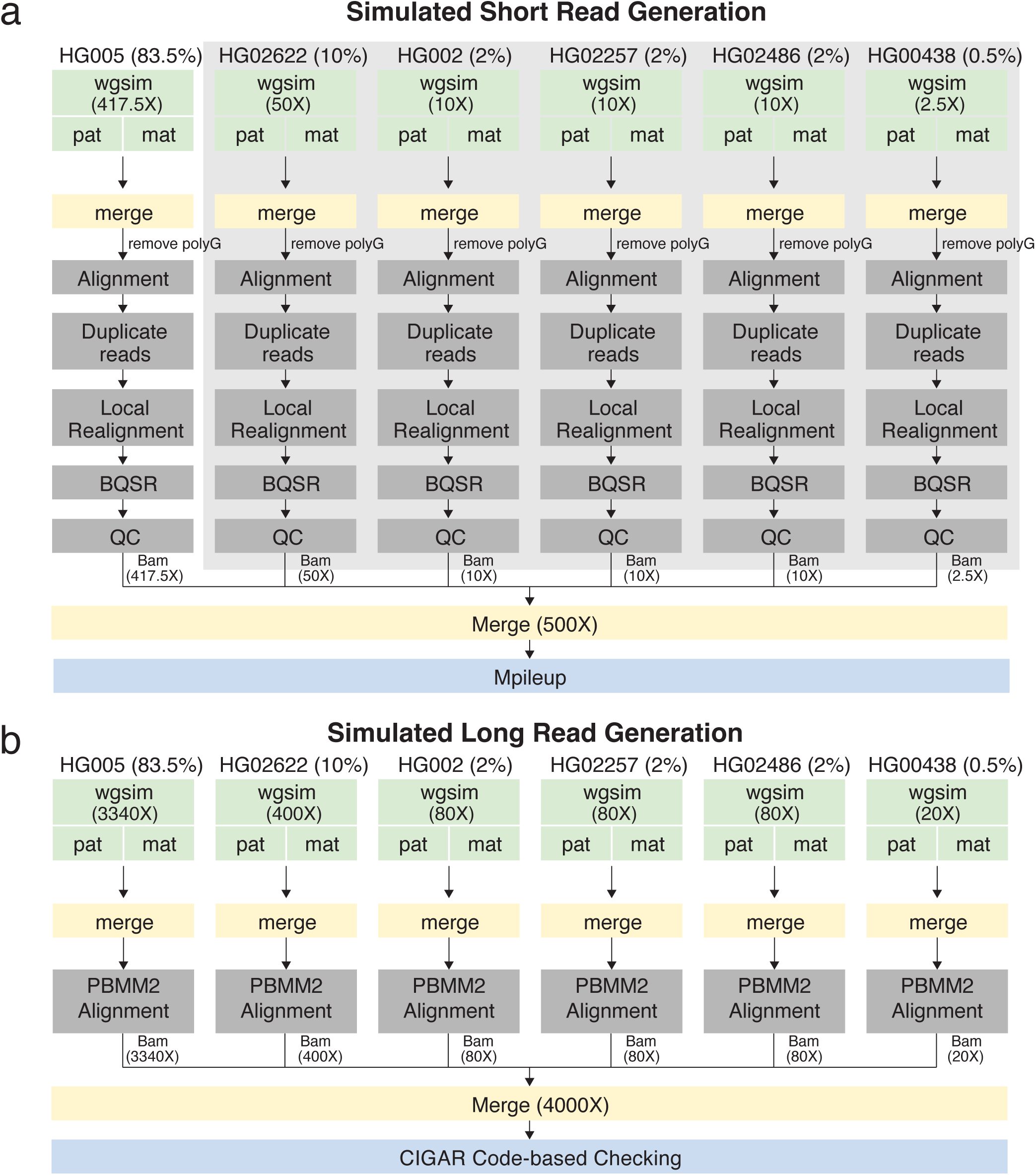
Simulated read generation pipeline for short-read and long-read sequencing data. (a) Detailed workflow showing the process of generating simulated short based on the haplotype assemblies for the six HapMap samples. At the determined mixing ratios, each sample undergoes initial processing steps including alignment, duplicate removal, local realignment, and quality control, followed by final merging at 500× coverage and mpileup. (b) Workflow showing the process of generating simulated long read. Simulated long-read data undergo alignment using “pbmm2”, followed by merging to reach a final coverage of 4000x and evaluation with CIGAR code checker.

**Figure S16.**
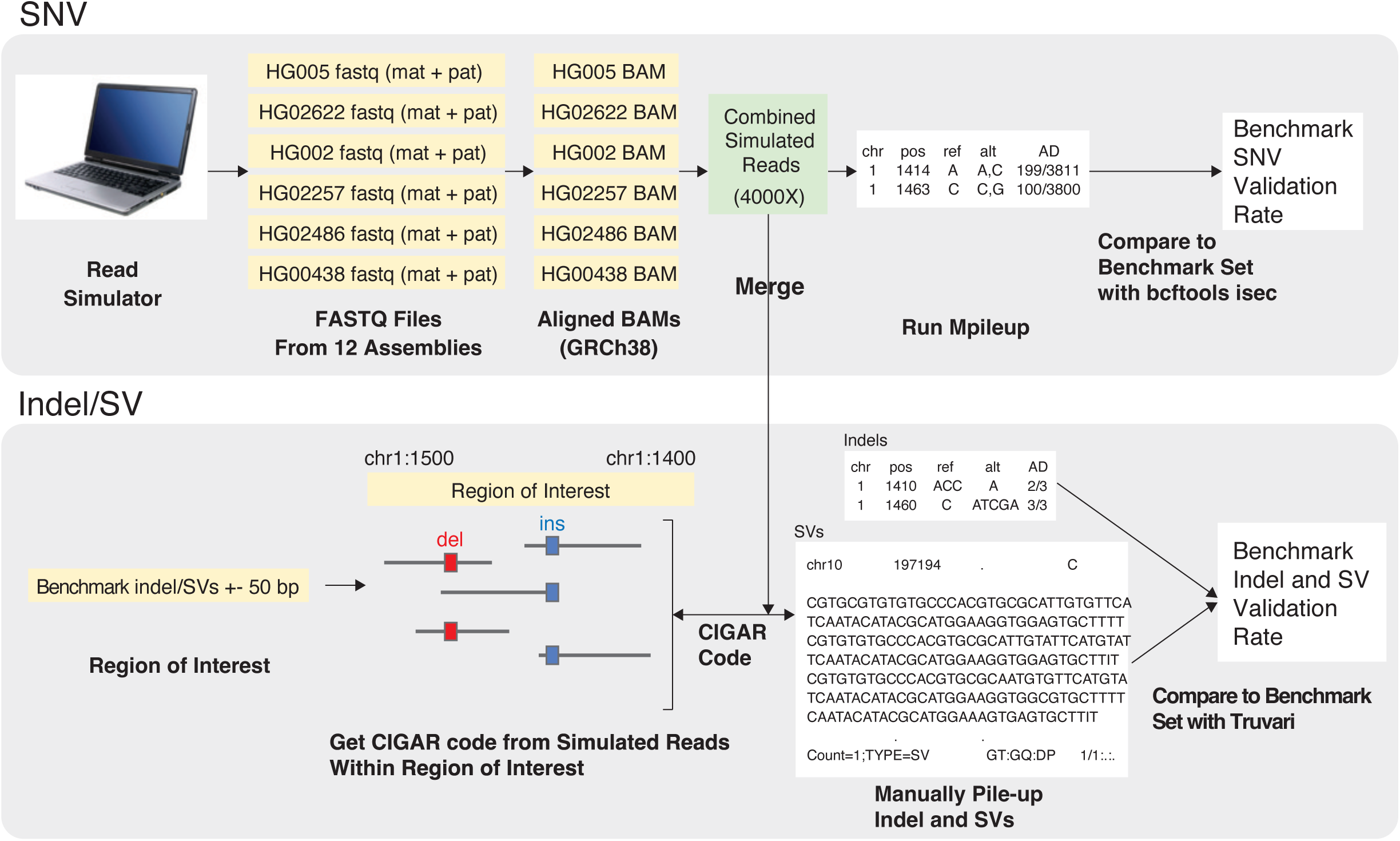
Benchmarking set validation strategy. Top: SNV validation workflow using wgsim simulator to generate reads from individual assemblies, followed by alignment and mpileup variant calling. Bottom: indel and SV validation approach based on CIGAR strings of reads within regions of interest, with validation rates shown for different variant types. SNV, single nucleotide variant; SV, structural variant.

